# Host community activity, but not always composition, explains viral biogeography in bulk and rhizosphere soils over a tomato growing season

**DOI:** 10.64898/2026.03.24.714046

**Authors:** Lucie Stern, Anneliek M. ter Horst, Konner E. Simpson-Johnson, Amélie C.M. Gaudin, Joanne B. Emerson

## Abstract

The soil microbiome is key to plant health and nutrient acquisition, and viruses likely play important but largely unknown roles in these processes. To interrogate bulk and rhizosphere soil viral biogeography, we collected samples over a tomato growing season in California from an experiment testing arbuscular mycorrhizal fungi (AMF) treatment. We generated 78 viromes, 16S rRNA gene, and ITS1 amplicon datasets, and 33 rhizosphere metatranscriptomes. Of 67,038 DNA viral ‘species’ genomes (vOTUs), 25% were previously identified, predominantely in agricultural systems, suggesting habitat filtering and greater viral homogeneity across agricultural compared to natural soils globally. Rhizospheres had significantly higher DNA viral richness than bulk soils, whereas no significant richness differences were observed for other biota. 60% of vOTUs were shared between compartments, compared to only 21-23% of bacterial and fungal taxa. Although bulk soil viral biogeography resembled that of prokaryotes, with significant structuring by moisture content, greater virome similarity between high-moisture bulk soils and rhizospheres suggests that conditions with high host activity selected for similar viral communities. In rhizospheres, while bacterial and fungal communities differed most over time, DNA and RNA viral communities differed most by sampling location, matching prokaryotic transcriptional patterns and further implicating host activity in viral biogeography. Similarly, AMF treatment induced changes in the prokaryotic transcriptome but, across biota, only significantly affected DNA viral communities. Overall, results indicate strong viral responses to spatiotemporally localized conditions, with viral biogeography reflecting both dispersal opportunities (high between neighboring bulk and rhizosphere soils, low across fields) and selection via local host activity.

## Introduction

Soils appear to host the most diverse and dynamic viral communities globally (Fierer, 2017), yet the ecological forces structuring these communities across soil compartments, space, time, and management remain largely understudied. Soil microbes contribute to essential agroecosystem processes, such as nutrient cycling and soil fertility, and rhizospheres are hotspots of microbial activity and plant-microbe interactions (Edwards et al., 2015; Mendes et al., 2013; Pieterse et al., 2016; Trivedi et al., 2020). Through root exudation, plants feed carbon-rich substrates to rhizosphere-associated microbial communities, affecting their activity and composition (Edwards et al., 2015; Mendes et al., 2013; Pieterse et al., 2016; Trivedi et al., 2020). In return, beneficial soil microbes help plants resist diseases, absorb nutrients, and produce hormones that promote growth (Bakker et al., 2018; Bulgarelli et al., 2013; Das et al., 2022; Edwards et al., 2015; Mendes et al., 2011; Pieterse et al., 2016). While agriculture-associated bacteria and fungi are relatively well studied, the role of agricultural soil viruses remains comparatively overlooked, despite a number of reviews highlighting their potential roles in microbial mortality and soil and plant health (Banerjee & van der Heijden, 2023; Emerson, 2019; Kimura et al., 2008; Pratama & Elsas, 2018; Williamson et al., 2017).

Compositional differences between bulk and rhizosphere soils are well established for bacterial and fungal communities but remain largely unexplored for soil viruses. The rhizosphere, defined as the narrow zone of soil influenced by root activity, differs noticeably from bulk soil in its physicochemical properties, including nutrient availability, pH, redox conditions, and gas exchange, largely as a result of root exudation and intense plant-microbe interactions (Bi et al., 2021; Hinsinger et al., 2005; Mendes et al., 2013; Williams & de Vries, 2020). These differing environmental conditions strongly structure microbial communities between soil compartments, often selecting a subset of taxa from the surrounding bulk soil and modifying microbial diversity and activity near roots (Bakker et al., 2015; Bakker et al., 2013; Ling et al., 2022). Although far fewer studies have explicitly studied soil viral communities between these compartments, available evidence suggests that viral communities also differ between bulk and rhizosphere soils (Swanson et al., 2009; Pratama & Elsas, 2018; Bi et al., 2021), potentially reflecting both shifts in host composition and changes in soil chemistry. In bulk soils, viral community composition has been linked to environmental factors, such as pH, moisture, depth, altitude, and other soil chemistry across diverse ecosystems (Adriaenssens et al., 2017; Bi et al., 2021; Emerson et al., 2018; Lee et al., 2022; Santos-Medellín et al., 2022; ter Horst et al., 2021; R. Wu, Davison, Gao, et al., 2021).

However, because the rhizosphere represents a zone of increased microbial activity and relatively buffered microenvironmental conditions compared to bulk soils, particularly with respect to soil moisture dynamics (Carminati et al., 2010, 2016; Benard et al., 2019), it may impose distinct selective pressures on viral communities. Whether soil viruses respond to rhizosphere conditions in ways similar to their presumed bacterial and fungal hosts remains to be investigated.

While early studies of soil viruses predated high-throughput sequencing (Fierer et al., 2007; Williamson et al., 2005) or relied largely on bioinformatic mining of DNA viral sequences from total metagenomes (Emerson et al., 2018; Paez-Espino et al., 2016; Trubl et al., 2018), recent advances have enabled more comprehensive investigations of soil viral communities (Nicolas et al., 2023; Santos-Medellin et al., 2021; Sorensen et al., 2021; ter Horst et al., 2021; Trubl et al., 2016, 2019). By purifying the viral fraction through 0.22 μm filtration, followed by library construction from comparatively low-input DNA (Emerson, 2019), it is now possible to recover a much greater DNA viral diversity from each soil sample (Santos-Medellin et al., 2021; ter Horst et al., 2021). By leveraging the conserved RNA-dependent RNA polymerase (RdRp) protein sequence, the diversity of viruses with RNA genomes can also be comprehensively assessed from RNA sequencing data (Hillary et al., 2022; Starr et al., 2019). Here we use both DNA viromics and, from rhizospheres, also metatranscriptomics to explore DNA and RNA viral biogeography in agricultural systems.

Recent DNA viromic studies have shown that soil viral communities commonly display strong spatial and temporal structuring, particularly in natural soils (Fudyma et al., 2025). Natural soil viral communities differ over meter-scale distances, and significant differences among habitats have been demonstrated at both regional and global scales (Durham et al., 2022; Fudyma et al., 2024; Santos-Medellín et al., 2022; ter Horst et al., 2021; R. Wu, Davison, Nelson, et al., 2021). Globally, substantial diversity and under-sampling have been indicated for viruses from a variety of natural soils, with only 1.5 to 4% of local viral ‘species’ (vOTUs) previously detected elsewhere (Geonczy, ter Horst, et al., 2025; ter Horst, Fudyma, Sones, et al., 2023). Natural soil viral communities also tend to display significantly higher rates of temporal succession than their prokaryotic counterparts (Santos-Medellín et al., 2023). Although hints of local spatiotemporal dynamics have been indicated for some agricultural soil viral communities (Santos-Medellin et al., 2021) results have been inconsistent (Sorensen et al., 2023) and derived from small sample sets (16-18 viromes) that were not specifically designed to measure spatiotemporal patterns. Thus, the extent to which strong spatial and temporal soil viral biogeographical patterns and potential decoupling from host biogeography extend to managed agricultural soils and/or rhizospheres remains poorly understood.

There is growing interest in using microbial amendments to improve soil health and crop productivity (Chaparro et al., 2012; Nuzzo et al., 2020), yet their consequences for soil microbiomes are largely unknown (Turina et al., 2018). AMF have received considerable attention as potential plant-beneficial amendments because of their widespread symbiotic associations with plants and their roles in nutrient acquisition and stress tolerance (Gosling et al., 2006; Guzman et al., 2021). AMF are increasingly used as bio-inoculants to enhance plant growth and resilience under abiotic stresses (Marro et al., 2022, Balliu et al., 2015; Hajiboland et al., 2010). However, in intensive cropping systems, such as modern tomato production, AMF benefits are often inconsistent and strongly context-dependent, varying with soil fertility, water availability, plant genotype, and management practices (Balliu et al., 2015; Chen et al., 2025; Wan et al., 2025; S. Wu et al., 2022). Thus, how AMF application impacts soil viral communities warrants further investigation to better understand how viruses might influence microbial community dynamics in response to these bio-inoculants. More fundamentally, a field-based AMF treatment experiment provides a testable framework for assessing short-term habitat filtration impacts on viral and microbial assembly.

Overall, despite growing interest in understanding how soil viruses influence plant-associated microbiomes in agroecosystems, we still know little about the ecological drivers shaping viral communities in agricultural soils or how these dynamics compare to those of bacteria and fungi. To address this gap, we generated 78 viral size-fraction metagenomes (viromes) to characterize viral community composition in 36 tomato plant rhizospheres and their associated bulk soils (plus 6 bulk soils before planting) throughout one growing season in an agricultural field in Davis, CA, USA. We used 33 rhizosphere metatranscriptomes (of 36 attempted) to characterize RNA viral communities and microbial gene expression, and we also generated 78 16S rRNA gene and ITS1 amplicon sequencing datasets from the same samples as viromes to investigate putative host bacterial and fungal communities. Half of the tomato plants were treated with AMF. Considering soil compartment, plot location, plant growth stage (time point), soil chemistry, and AMF treatment, we evaluated the extent to which viral biogeography reflected expected patterns from other microbiota. We also leveraged the PIGEONv2.0 database of 443,140 vOTUs (ter Horst, Fudyma, Sones, et al., 2023) to explore regional-to-global biogeographical distributions of vOTUs in our study.

## Results and Discussion

### 1. Dataset overview

A total of 67,038 viral operational taxonomic units [vOTUs, approximately species-level taxonomy; (Roux et al., 2019)] was recovered from the 78 viromes, 2% of which could be taxonomically classified [via vConTACT2, (Bin Jang et al., 2019)]. We recovered 380 viral RNA-dependent RNA polymerase (RdRp) genes (Hillary et al., 2022; Starr et al., 2019; ter Horst, Fudyma, Bak, et al., 2023) from the 33 rhizosphere metatranscriptomes, which were dominated by predicted viruses of fungi and prokaryotes. Specifically, there were 174 putative viruses of fungi, 166 of prokaryotes, 20 of insects and/or vertebrates, 15 of other animals and/or plants, and only 3 confidently assigned to plants alone. A total of 1,051,814 transcripts were recovered from the metatranscriptome data, and ∼30% were functionally annotated, of which 96% were bacterial, <1% archaeal, < 1% plants, < 1% fungal, ∼3% other eukaryotes, and <0.5% viral. As such, in downstream analyses, we interpret community-wide gene expression data from the metatranscriptomes as reflective of bacterial transcription. A total of 16,146 16S rRNA gene and 6,684 ITS1 amplicon sequence variants (ASVs) were obtained from the same 78 samples as the DNA viromes.

### 2. Agricultural soil viral ‘species’ (vOTUs) show more homogeneous detection across similar habitats than do those from natural soils

To investigate the global habitat distribution of vOTUs recovered at our field site (Russell Ranch), we leveraged the PIGEONv2.0 database (ter Horst, Fudyma, Sones, et al., 2023) of 443,140 vOTUs from a variety of ecosystems (Figure 1). Of the 67,038 vOTUs recovered in this study, 16,452 (25%) were from PIGEONv2.0 and thus previously detected in other studies, including 10,173 (15%) from the same field site in 2018 (our samples here were collected in 2021), suggesting that part of the virosphere at this site is stable and/or recurring over time. This is consistent with a previous observation of relative temporal stability over months for soil viral communities at this field site (60% of vOTUs shared between July and October of the same year, Sorensen et al., 2023) and indicates that viral populations may persist over long periods of time in agricultural systems.

**Figure 1.**
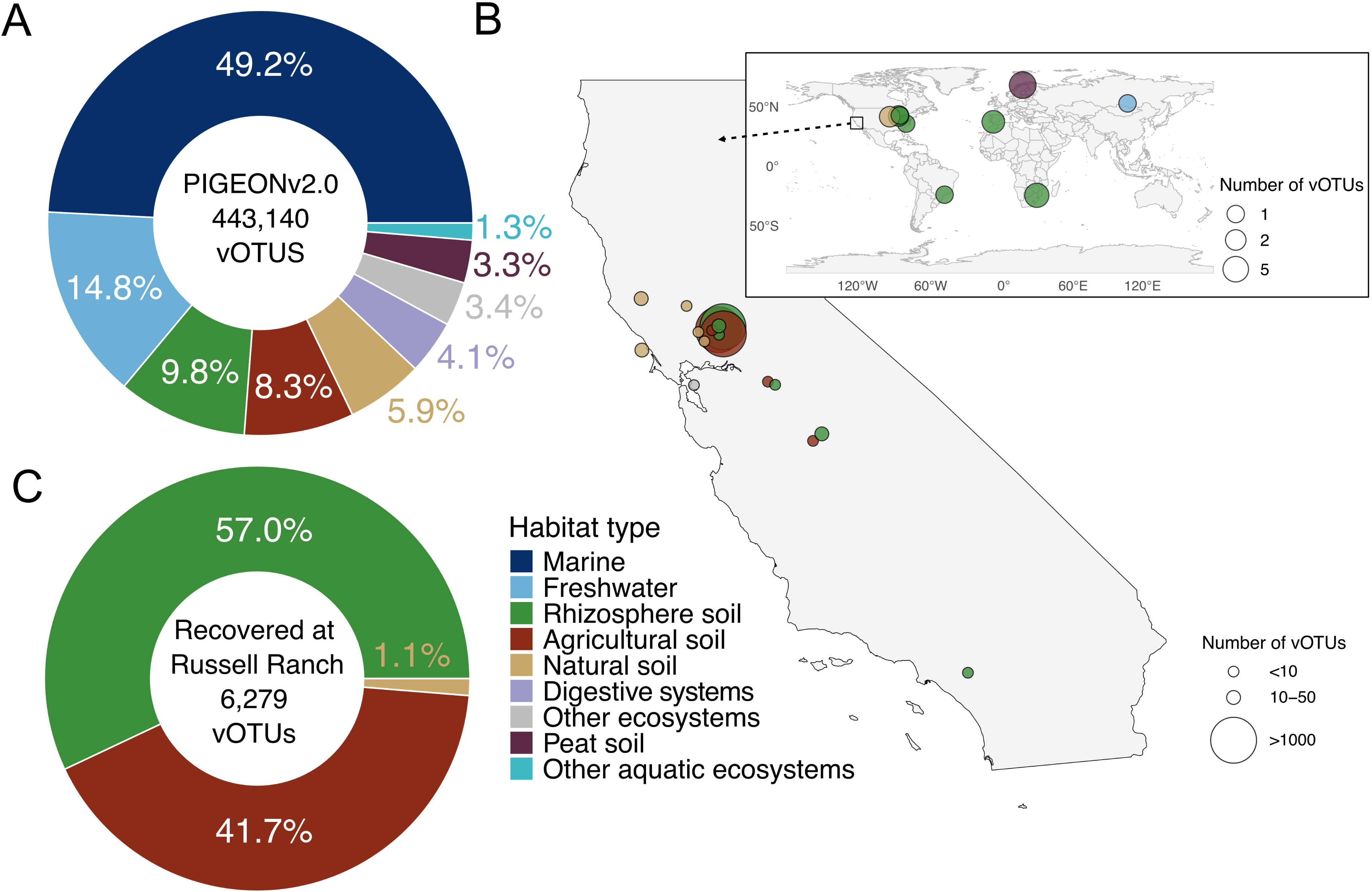
Global and California-wide distribution of Russell Ranch vOTUs, recovered using the PIGEON v2.0 database. A: Composition of the PIGEONv2.0 database of 443,140 vOTU sequences, colored by environment, excluding vOTUs from Russell Ranch. B: Global and California-wide locations of vOTUs (n = 6,279) recovered at Russell Ranch by read mapping to PIGEONv2.0 based on where they were first identified. C: Relative proportions of vOTUs (n=6,279) recovered at Russell Ranch by read mapping to PIGEONv2.0, grouped and colored by the environment from which each vOTU was first recovered;-0.2°/o of vOTUs are not depicted in C (slices too small to visualize, 0.1°/o peat, 0.02°/o freshwater, 0.06°/o other ecosystems) but appear in B.

After excluding all previously detected vOTUs from Russell Ranch (the same site as this study), the remaining 6,279 vOTUs (11% of the full dataset) detected here from PIGEONv2.0 were predominantly recovered from other agricultural sites, including both bulk and rhizosphere soil (Figure 1B and 1C). Of those, 98% were found in other agricultural systems in California (Figure 1B), with most of those (77%) found in bulk or rhizosphere soils from almond orchards (ter Horst, Adebiyi, Hernandez, et al., 2023). Extensive detection in California almost certainly reflects geographic sampling bias, as most (90%) of the soil vOTUs in PIGEONv2.0 are from our group’s locally sampled viromes in California, and soil viromes have not been extensively reported from other locations. Still, vOTU recovery here from predominantly agricultural systems suggests that these viruses are adapted to these habitats and/or their host communities, as only 18% of the vOTUs in PIGEONv2.0 are from agricultural or rhizosphere soils. More generally, vOTUs from our dataset were previously recovered in other agricultural bulk soils (n=2,619), other agricultural rhizosphere soils (n=3,580), natural soils (n=69), peat (n=6), and other ecosystems (n=4, compost; Figure 1C). Consistent with strong habitat boundaries, no vOTUs from marine ecosystems were recovered here, and only one vOTU was recovered from freshwater, whereas in wetlands, 46% of the previously detected vOTUs were from marine or freshwater environments (ter Horst, Fudyma, Sones, et al., 2023).

Among vOTUs originally described from rhizosphere studies, most (53%) were detected in both rhizosphere and bulk soils here, indicating substantial compositional overlap between compartments. Similarly, most (64%) vOTUs previously detected from agricultural soils were shared between rhizospheres and bulk soils here. These patterns suggest that many agricultural soil viruses are compartment generalists, and that, overall, agricultural soil and rhizosphere viruses exhibit distribution patterns consistent with agriculture-specific habitat filtering. Habitat specificity for soil vOTUs has been shown previously for peat, fresh and saline wetlands, and burned soils (Geonczy, ter Horst, et al., 2025; ter Horst et al., 2021; ter Horst, Fudyma, Sones, et al., 2023), and here we extend this likely habitat-specific selection (presumably at least partly by way of hosts) to agricultural soils and rhizospheres.

Though more sampling across soil habitats and locations is needed for broad generalizability, with 15% of vOTUs here previously detected (excluding those from the same field site), relative to only 1.5 to 4% in prior studies of natural systems (Geonczy, Hillary, et al., 2025; ter Horst et al., 2021; ter Horst, Fudyma, Sones, et al., 2023), agricultural soil viral ‘species’ seem to have comparatively more homogeneous and stable regional-to-global distributions than those in natural soils. Although sampling bias is clearly a contributing factor, prior analyses of natural soils suffered from similar biases (ter Horst et al., 2021; ter Horst, Fudyma, Sones, et al., 2023); in both types of soils (natural and agricultural), most previously detected vOTUs were from similar habitats that had undergone prior viral sequencing and/or specific viral analysis efforts. Together, these results show that the same agricultural soil viral ‘species’ (vOTU) can be detected over large spatial distances, including on multiple continents, that habitat characteristics play an important role in shaping agricultural soil and rhizosphere viromes, and that agricultural soil viruses seem to be more uniformly distributed over space and/or time than those in natural systems.

### 3. Viral communities in bulk and rhizosphere soils diverged over the course of the tomato growing season

To interrogate the main drivers of community structure in our dataset, we first explored which variables shaped the viral, bacterial, and fungal communities overall, across all time points, locations, compartments (bulk vs. rhizosphere), and AMF treatment. Overarching patterns were similar across biota, with time, compartment, and plot (location) all contributing factors for DNA viral, bacterial, and fungal communities, followed by significant but marginal effects of AMF treatment, as well as contributions from interactions among variables (Table S1-S3). However, the proportions of variance explained (R^2^ values) were much higher for viruses (R^2^ = 0.158, 0.152, 0.135 for time, compartment, and location, respectively), with 80% of the total variation in DNA viral communities explained by these factors or their combination (Figure 2A and Table S2). By comparison, only time substantially explained variance for bacterial communities (R^2^ = 0.117, still lower than the R^2^ for any of the top three factors for viruses), and all other R^2^ values for correlations with bacteria and fungi were less than 0.1, with only 60% of the total variance explained by these factors for bacterial and fungal communities (Figure S1 and Table S2). Mantel tests revealed a strong correlation between bacterial and fungal community beta-diversity patterns (R = 0.67, p = 0.001) and a significant but less strong correlation between DNA viral and bacterial community beta-diversity (R = 0.33, p = 0.001; Table S4). Together, these results indicate that, although there is evidence for viral responses to host community composition, viral communities more strongly reflected spatiotemporally localized conditions (as measured across multiple variables) than did bacterial or fungal communities, such that host community composition alone did not fully explain viral biogeography.

**Figure 2.**
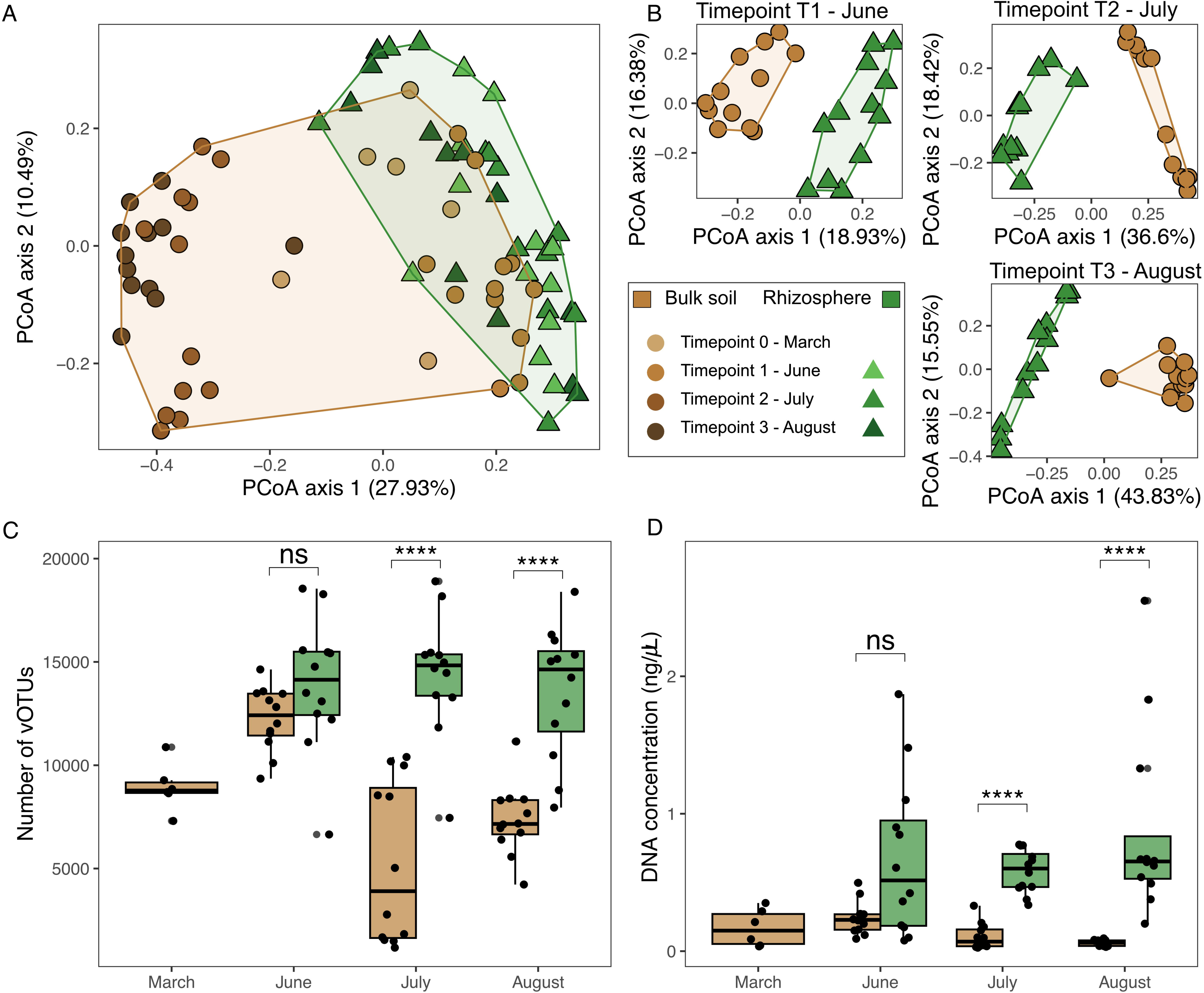
Bulk and rhizosphere soil viral community composition, viral’species’ (vOTU) richness, and viromic DNA concentrations. (A) Principal coordinates analysis (PCoA) of DNA viral communities across 78 viromes, based on Bray-Curtis dissimilarities of vOTU relative abundances. Points are colored by soil compartment (bulk soil vs. rhizosphere) with shades representing sampling time points: TO (March, pre-planting), T1 (June, 6 weeks after planting), T2 (July, 10 weeks after planting), and T3 (August, 17 weeks after planting), according to the panel B legend. Axis labels indicate the percentage of total variance explained. (B) Separate PCoAs of DNA viral communities, faceted by sampling time point and colored by soil compartment. (C, D) Boxplots of (C) total vOTU richness (number of vOTUs) per virome per time point in bulk and rhizosphere soils and (D) DNA concentrations (a proxy for viral particle abundances) obtained from virome extractions from bulk and rhizosphere soils across sampling time points, colored by soil compartment, as in (B). Boxes represent the interquartile range (IQR, 25th-75th percentile), the central line shows the median, whiskers extend to 1.Sx IQR, and individual points outside whiskers are outliers. Points are jittered for clarity. Statistical significance between bulk and rhizosphere vOTU richness (C) and DNA concentration (D) was assessed using a Wilcoxon rank-sum test and is indicated where applicable: ns., not significant; *p < 0.05; **p < 0.01; ***p < 0.001; ****p< 0.0001.

Although DNA viral community composition differed significantly between bulk and rhizosphere soils, the strongest compositional separation for viruses was between later season bulk soils from July and August and all of the other viromes (March and June bulk soils and all rhizospheres together; Figures 2AB). Viral communities showed greater compositional overlap between bulk and rhizosphere soils early in the season, followed by increasing divergence later in the growing season, a pattern consistent with but less pronounced in bacterial and fungal communities (Figure 2B and S1CD). Overall, 60% of vOTUs were shared between compartments, compared to only 21% and 23% of bacterial and fungal ASVs, respectively (Figure S2). There were no significant differences in richness between compartments for bacteria or fungi (Figure S1EF), but viral richness was significantly higher in rhizospheres at the last two time points, with bulk soil viral richness decreasing at later time points, while rhizosphere viral richness remained relatively stable and high (Figure 2C). This pattern was mirrored by viromic DNA yields (an established proxy for viral biomass, Santos-Medellín et al., 2023), with significantly higher viral biomass in rhizospheres compared to bulk soils at the last two time points (Figure 2D). Prior mining of viral sequences from total metagenomes from late-season maize rhizosphere and associated bulk soil samples reported strong soil compartmental differentiation of soil viral communities (Bi et al., 2021), and our results are consistent with that finding but suggest that, earlier in the growing season, these compartmental differences may not be as pronounced. Still, with some notable differences (more shared viral ‘species’ between compartments and significantly higher viral richness in rhizospheres), the overall biogeographical patterns for soil and rhizosphere DNA viruses resembled those of their likely host prokaryotes.

### 4. Bulk soil viral communities responded significantly to changes in soil moisture, likely indirectly via host community responses and host activity

We next sought to explore the relative importance of time, plot location, and AMF treatment (key experimental design factors that could be reflective of succession, dispersal limitation, and environmental selection, respectively) in structuring bulk soil viral communities. These communities were primarily structured by time point (R^2^= 0.414, P = 0.001; Figure 3A and 3B) and secondarily by plot location (R^2^= 0.149, P = 0.001; Figure S3A), followed by a weaker effect of AMF treatment (R^2^= 0.026, P = 0.007; Figure S3B), as well as interactions among variables (Table S5). Bulk soil bacterial and fungal communities showed similar patterns, being structured mainly by time and plot location, with no significant effect of AMF treatment (Table S5, Figure S3C-F). Thus, at a coarse scale, bulk soil DNA viral communities appeared to be following the behavior of their likely host bacteria (Table S3).

**Figure 3.**
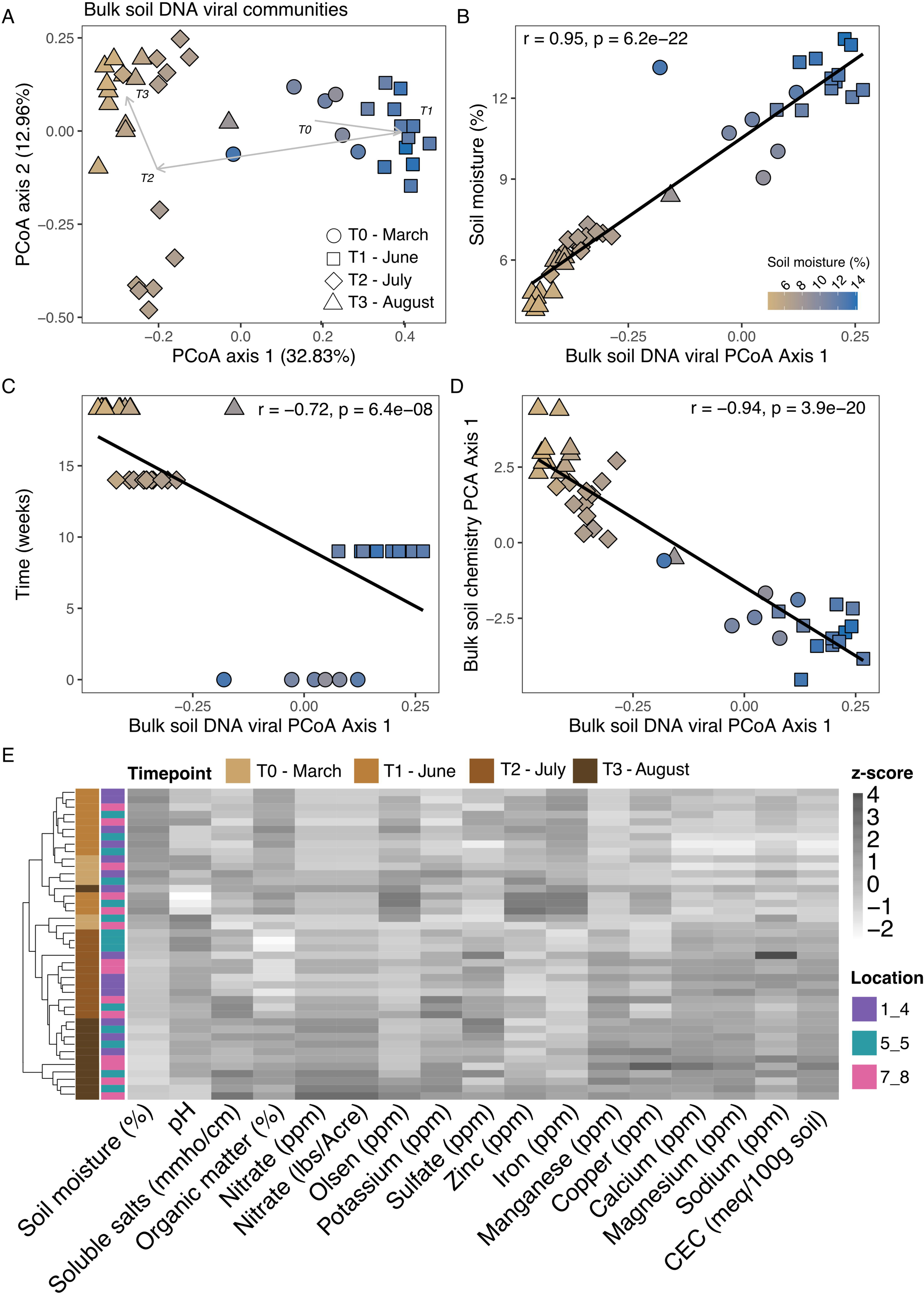
Bulk soil DNA viral community composition over time, moisture content, and other soil physicochemical properties. (A) Principal Coordinates Analysis (PCoA) of DNA viral community composition, based on Bray-Curtis dissimilarities of vOTU relative abundances, colored by bulk soil moisture content (according to the legend in B), with shapes indicating time point. (B-C) Linear regression analyses, comparing PCoA 1 of Panel A (a proxy for bulk soil DNA viral community composition) to (B) soil moisture content, (C) time of sample collection, and (D) a proxy for overall physicochemistry: the first axis of a principal component analysis (PCA axis 1, plot in Fig S3G) of all bulk soil physicochemistry measurements (shown in E), excluding soil moisture, with correlation statistics for (B-C) inset in each graph. (E) Hierarchical clustering of samples, based on z-scores of measured bulk soil physicochemical variables. mmho/cm, millimhos per centimeter; ppm, parts per million; CEC, cation exchange capacity; meq/100 g, milliequivalents per 100 grams soil

The observed compositional differences among time points raised the question of whether temporal changes reflected a directional successional trajectory of community turnover or instead consisted of discrete, non-directional shifts, potentially driven by selection imposed by other variables, such as changing physicochemical conditions. We thus tested whether treating time as a linear variable (weeks since the start of sampling) versus as discrete categorical sampling timepoints changed the influence of time on community composition (Table S6). Categorical timepoints explained approximately twice as much variation in the DNA viral communities as did linear time (with bacterial and fungal communities showing similar or slightly stronger effects), suggesting that the significant influence of time point was likely due to fluctuations in environmental conditions and associated selective pressures (by way of hosts), as opposed to iterative temporal turnover.

The observed differences in bulk soil viral communities across time points, including divergence from rhizospheres at the last two time points, coincided with lower soil moisture towards the end of the growing season. Moisture dropped from an average of 12% in March and June to 6% in July and August and a PCoA plot of bulk soil viral community composition revealed an apparent separation by soil moisture along the PCoA axis 1 (Fig 3A). Indeed, in a linear regression, the PCoA axis 1 for bulk soil viral communities was strongly and significantly correlated with soil moisture (R = 0.95, p < 0.001; Figure 3B) and was still significantly but less strongly correlated with linear time (R =-0.72, p < 0.001; Figure 3C). Comparable correlations with soil moisture were observed for bacterial PCoA axis 1 (R = 0.84, p < 0.001; Figure S4A) and fungal PCoA axis 2 (R = 0.79, p < 0.001; Figure S4B), likely indicating an indirect viral response to soil moisture by way of more direct host responses. However, while viral richness (Figure S4C; R = 0.43, p < 0.001) and viromic DNA yields as a proxy for viral biomass (Figure S4D; R = 0.34, p < 0.001) were also positively correlated with soil moisture, no comparable relationship emerged for bacterial or fungal richness (Figure S4E and S4F). Together, these patterns suggest a strong dependence of DNA viruses and their presumed hosts on soil moisture, consistent with previous studies showing that moisture regulates viral abundance, persistence, and mobilization in soils (Cao et al., 2022; Santos-Medellín et al., 2023; Williamson et al., 2017; R. Wu, Davison, Gao, et al., 2021; R. Wu, Davison, Nelson, et al., 2021).

Although soil moisture emerged as a clear explanatory factor for bulk soil viral community composition, the relative contributions of host community composition and other soil chemistry variables required further investigation. All microbial groups showed significant correlations with one another in bulk soils, with the strongest associations observed between bacterial and fungal communities (Mantel R = 0.72, p = 0.001) and, for DNA viruses, with bacterial communities (Mantel R = 0.57, p = 0.001, Table S4). In bulk soils, viral communities showed the strongest correlations with soil physicochemical properties (Mantel R = 0.59, p = 0.001, Figure 3D), whereas bacterial communities were significantly but more weakly correlated (R = 0.33, p = 0.001), and fungal communities showed no significant correlation with soil chemistry (p = 0.138; Table S4). While soil chemical properties (without soil moisture included) overall were highly significantly correlated with bulk soil viral community composition (R =-0.94, p < 0.001; Figure 3D), the correlation was just as strong for soil moisture alone (R =-0.93, p <0.001). In Figure 3, a sample from timepoint T3 clustered with earlier timepoint samples rather than other T3 samples across all ordinations, driven by divergent soil chemistry than other samples from the same timepoint (Figure 3E).

In a variance partitioning analysis, where soil moisture was considered separately from other soil chemical variables; among these, only pH and nitrate were retained as significant predictors of DNA viral communities (Table S7). This is consistent with previous work showing that pH and soil chemical composition can influence viral attachment, persistence, and virus-host encounter and infection rates in soils (Chariou et al., 2019; Gerba, 1984; Loveland et al., 1996; Williamson et al., 2017). Variance partitioning also revealed strong covariance among time point, soil moisture, and soil chemistry, explaining why categorical time points captured most of the variation in bulk soil DNA viral communities, whereas linear temporal progression did not.

Together, these results may also explain why bulk soil viral communities were most similar to rhizospheres early in the growing season: soils with comparatively high moisture content and rhizospheres are both known hotspots of microbial activity. Consistent with greater viral production and diversity in rhizospheres and under high moisture content in bulk soils, conditions known to promote the activity of other microbes(Bakker et al., 2013; Brockett et al., 2012; E. Evans et al., 2022; Y. Li et al., 2017; Raaijmakers et al., 2009; Reinhold-Hurek et al., 2015), we hypothesize that higher soil moisture early in the season supported greater host activity and viral mobility (dispersal) between bulk and rhizosphere soils, resulting in more homogeneous viral communities between compartments. Soil moisture is already a well-established driver of soil viral community structure (Santos-Medellín et al., 2023; R. Wu, Davison, Gao, et al., 2021) and of microbial activity more broadly (Brockett et al., 2012; E. Evans et al., 2022; Y. Li et al., 2017). Previous work has shown that rhizosphere soils retain water during drying periods, while bulk soils experience greater moisture fluctuations (Benard et al., 2019; Carminati et al., 2016), so community responses to rhizosphere moisture would be expected to be less pronounced than in bulk soils. As further evidence for sustained moisture conditions within the rhizosphere throughout the growing season, rhizosphere metatranscriptome data showed no evidence of increasing desiccation or osmotic stress over time in control plants (p > 0.05, Table S8). DNA repair machinery, and oxidative stress responses, all of which are hallmarks of the bacterial desiccation response (Scales et al., 2023), indicates that rhizosphere microbiomes were not experiencing water limitation. This supports the concept of the rhizosphere as a buffered microenvironment where plant-derived water and carbon inputs maintain microbial activity even as bulk soil conditions fluctuate. Microbial activity has been hypothesized to be an important driver of soil viral community composition (Santos-Medellín et al., 2022, 2023), and here we suggest that microbial activity could be a stronger selective force in viral community assembly than the habitat-specific selection imposed by soil compartment.

### 5. Spatial structuring was the dominant feature of rhizosphere DNA and RNA viral communities, reflecting prokaryotic activity patterns more than community composition

Considering the overall experimental design features of time point, plot location, and AMF treatment, we next examined the ecological drivers of rhizosphere viral community structure to assess whether patterns observed in bulk soils also applied at the plant-root interface. Unlike bulk soils, which were far and away structured by time point (and associated moisture and chemistry differences), rhizosphere DNA viral communities were primarily structured by plot location (R² = 0.272, P = 0.001; Figure 4A), followed by AMF treatment (R² = 0.135, P = 0.001, Figure 4B), with a smaller but still significant effect of sampling timepoint (R² = 0.079, P = 0.003; Table S9). Similar spatial structuring was observed for rhizosphere RNA viral communities (Figure 4C), which also differed significantly by plot location (R² = 0.103, P = 0.001), timepoint (R² = 0.077, P = 0.005), and their interaction (R² = 0.133, P = 0.034; Table S9). These compositional differences among locations at the same field site are consistent with previous reports of strong spatial structuring of natural soil viral communities at local scales (Durham et al., 2022; Santos-Medellin et al., 2022) and suggest dispersal limitation as an important factor in rhizosphere viral community assembly.

**Figure 4.**
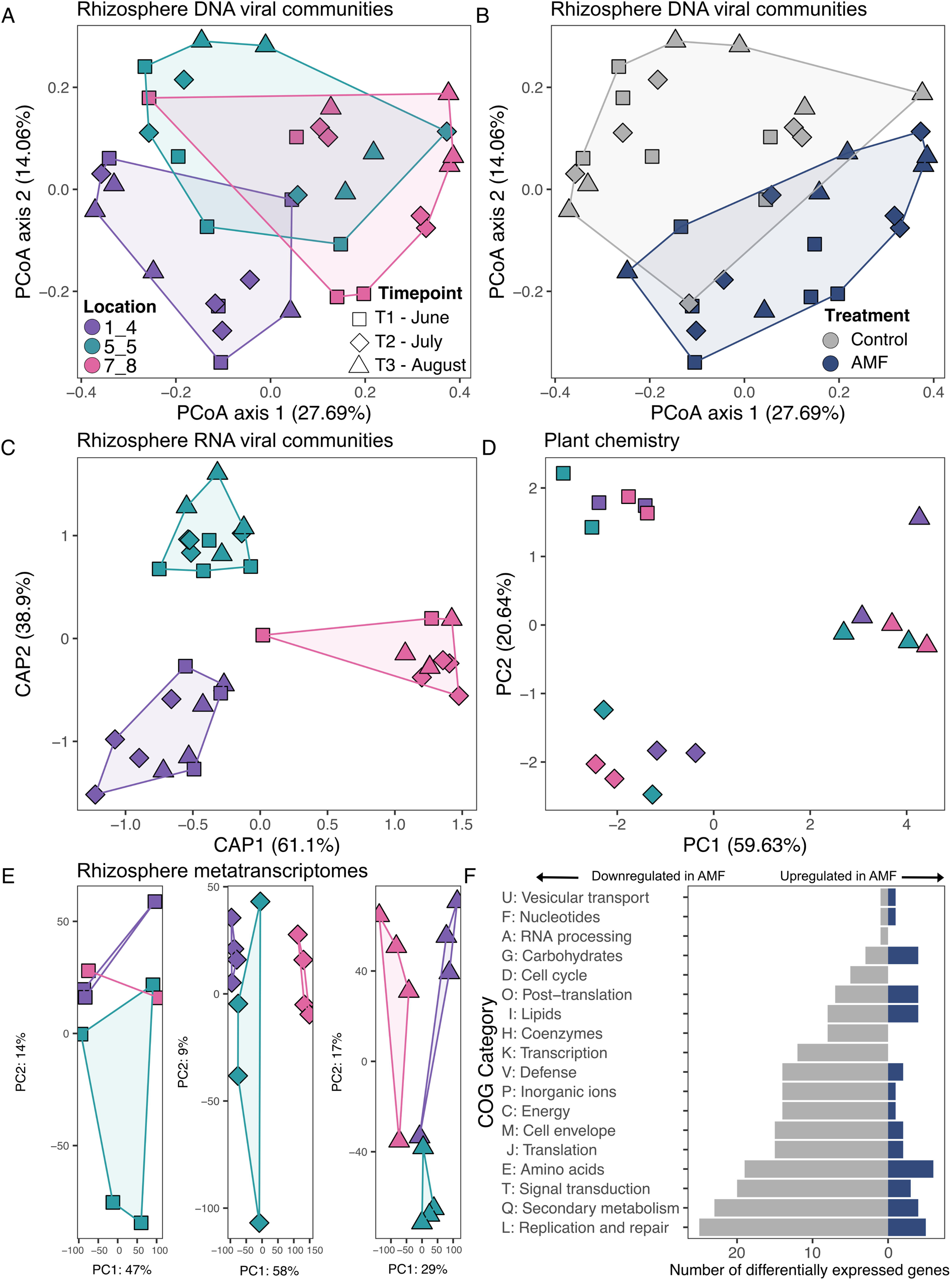
Rhizosphere DNA and RNA viral biogeography and explanatory factors. (A,B) Principal Coordinates Analysis (PCoA) of rhizosphere DNA viral community composition, based on Bray-Curtis dissimilarities of vOTU relative abundances (same plot in A and B, colored by plot location in A and AMF treatment condition in B), (C) Constrained Analysis of Principal coordinates (CAP) of rhizosphere RNA viral communities, based on Bray-Curtis dissimilarities of RNA-dependent RNA Polymerase (RdRP) gene relative abundances, (D) Principal Components Analysis (PCA) of a suite of 13 z-scores plant aboveground chemistry measurements (Table S13), (E) PCA of rhizosphere metatranscriptomic data, based on read mapping to individual genes assembled from the metatranscriptomes (predominantly prokaryotic data, as >96% of the genes were predicted to be derived from prokaryotes). Each gene was treated as a separate feature, and gene relative abundances across samples were used to calculate Euclidean distances, (F) Number of differentially expressed prokaryotic genes between AMF treatment and control samples, grouped by COG category (full category descriptions provided in Table S14). Shapes in all ordination graphs (A-E) correspond to time point. Colors in A, C, D, and E correspond to plot location, according to the legend in A. Colors in B and F correspond to AMF treatment, according to the legend in B. For PCA and PCoA, percentage values indicate the proportion of total variance in the dataset explained by each axis. For CAP, axes are constrained to maximize separation between predefined groups, and percentages sum to 100°/o of the variance that can be attributed to group differences, not total variance in the dataset.

In contrast to the viral communities, rhizosphere bacterial and fungal communities were structured primarily by time point, likely reflecting shifts in plant growth stage and associated changes in root exudates (Liu et al., 2022; McLaughlin et al., 2023; Upadhyay et al., 2022), consistent with significant differences in plant chemistry over time but not location (Figure 4D, Table S9-S10). As rhizosphere bacterial communities from the same plant species can remain relatively uniform across broad geographic regions (Ling et al., 2022; Xu et al., 2018), compositional similarities within the same field were unsurprising, but the decoupled biogeographical patterns between communities of viruses and their likely hosts is a new observation in the rhizosphere. Despite these differing biogeographical patterns, Mantel tests confirmed that rhizosphere DNA viral community composition still correlated strongly with both prokaryotic and fungal communities (R = 0.35-0.37, p < 0.001), though prokaryotes and fungi were more strongly correlated with each other (R = 0.56, p < 0.001, Table S4). RNA viral communities showed no significant correlations with prokaryotes or fungi, likely due to the broad taxonomic heterogeneity of hosts for RNA viruses (all domains of life), though more granular tests of viral subgroups according to predicted host taxonomy still revealed no significant correlations with their host communities (Table S4). More consistent with the viral community spatial patterns, metatranscriptomic data (derived from transcript mapping to assembled metatranscriptomic data) were most strongly differentiated by plot location, followed by time point and AMF treatment (Figure 4E, Table S9), with divergence among sampling locations increasing later in the season. As the transcripts predominantly reflected prokaryotic activity, these results are consistent with host activity as a key driver of rhizosphere viral community composition, particularly for DNA viruses.

Overall, we suggest that the observed rhizosphere viral biogeographical patterns arose from multiple features of viruses and their biotic and abiotic requirements, many of which differ from those of their hosts. Spatial structuring indicates that dispersal limitation had a strong impact on rhizosphere viral community assembly, consistent with short residence times for viral particles (estimated to be on the scale of days in active soils; DiPietro et al., 2023; Santos-Medellín et al., 2023) and previous observations of spatial structuring for viruses in natural soils (Santos-Medellin et al., 2021). For a productive infection, a viral particle needs to first encounter (by way of dispersal) a suitable host cell, and then that host cell needs to be sufficiently physiologically active for an infection to proceed (Williamson et al., 2017). Temperate phages (those capable of lysogeny) similarly rely on host cell activity for lysis-lysogeny decisions (Williamson et al., 2017), such that both lytic and temperate phage responses to host activity could be captured in the viral particle fraction (as measured in viromes here) and reflected in biogeographical patterns. Altogether, the additional limitations and selective pressures on viruses, beyond those shaping bacterial and fungal communities, resulted in decoupled biogeography for rhizosphere DNA viruses and their host prokaryotic communities, with spatial structuring and host activity patterns best predicting rhizosphere viral community composition.

### 6. AMF treatment induced changes in bacterial transcription that correlated with changes in the rhizosphere DNA viral community

We next explored in more detail how AMF treatment influenced viral community composition, especially in the rhizosphere, where localized environmental conditions and host activity might play an especially strong role in shaping viral communities. Although AMF application (via root-dipping pre-planting) did not significantly affect root AMF colonization rates or a suite of measured plant traits at harvest (Figure S5; Table S10), AMF treatment did have a significant effect on DNA viral community composition in both bulk soils and the rhizosphere, with a strong effect in the rhizosphere and only a marginal effect in bulk soils (Figure 4B; Table S5, S9). In contrast, AMF treatment had no detectable effect on bulk soil bacterial or fungal community composition and only a marginal effect on those communities in the rhizosphere (Tables S5, S9). Similar conclusions have been drawn in other soil systems, where AMF alone exerted limited influences on microbial communities unless combined with additional organic amendments (Xiao et al., 2020). However, metatranscriptomic analyses revealed a transcriptional response to AMF application in the rhizosphere (Table S9, S11), with most transcripts of prokaryotic origin. Differential expression analysis (padj < 0.05, |log2FC| > 1; n = 496 genes) showed that transcripts across multiple functional categories (replication and repair, secondary metabolism, signal transduction) were broadly downregulated under AMF treatment (Figure 4F, Table S11), though most lacked functional annotation (∼40% were COG-annotated). Notably, a set of 22 desiccation and stress-related genes showed significantly increasing expression over time in AMF-treated plants but not in controls (Figure 4G, Table S8). This pattern resembles well-documented priming effect in AMF-colonized plants, where stress-responsive genes are upregulated even under non-stress conditions, enabling faster responses to subsequent stress (Mauch-Mani et al., 2017; Santander et al., 2017; Song et al., 2015) (Pozo et al., 2015; Santander et al., 2017). Our results suggest that a similar priming phenomenon may occur in the rhizosphere and indicate that AMF treatment influenced bacterial host physiology and activity more than community structure. As these functional changes co-occurred with shifts in DNA viral community composition, they provide additional evidence that DNA viral community composition was closely tied to host metabolic state and activity, rather than to host taxonomic composition alone.

## Conclusions

Here, we investigated viral community assembly and its ecological drivers in tomato bulk and rhizosphere soils over one growing season in California, USA, revealing strong ties between viral biogeography and both measured and presumed (previously established) patterns of prokaryotic activity. Agricultural soil viruses showed strict habitat boundaries, whereby nearly all previously known vOTUs were originally detected in other agricultural rather than natural ecosystems, implicating habitat filtering as a mechanism for maintaining stable or recurrent viral populations across agricultural ecosystems.

Viral communities were governed by distinct ecological processes compared to their microbial hosts. Expected differences between bulk and rhizosphere soils emerged for all measured microbial communities (prokaryotes, fungi, and viruses), but compositional separation between compartments increased over the growing season, most strongly for viruses. Bacterial and fungal communities responded primarily to temporal dynamics in both rhizosphere and bulk soil compartments, likely reflecting seasonal shifts in soil chemistry and plant phenology. In contrast, viral community assembly drivers were compartment-specific: bulk soil viral communities tracked temporal changes in moisture and chemistry, while rhizosphere viruses were structured by spatial location and AMF treatment. This decoupling between virus and host biogeographical patterns suggests that viruses experience stronger spatial constraints than do their larger hosts, challenging conventional assumptions about size-related dispersal in soils (Kuzyakov & Mason-Jones, 2018; Santos-Medellín et al., 2022). AMF treatment further showed decoupled patterns between virus and host communities, with significant restructuring of rhizosphere viral but not bacterial or fungal composition. However, AMF treatment induced widespread transcriptional shifts in the rhizosphere microbiome, including stress-gene priming, corresponding to previously documented plant responses to mycorrhizal colonization. This suggests that viruses track host metabolic states rather than taxonomic composition, responding to functional changes that remain invisible in standard community profiling. Together, these findings position soil viruses as dynamically responsive yet spatially constrained components of the soil microbiome, governed by ecological rules distinct from those shaping bacterial and fungal communities, and they underscore the need to integrate virus-specific ecology into broader frameworks for understanding soil microbial dynamics.

## Material and methods

### Sample collection and processing

Samples were collected at the Russell Ranch Sustainable Agriculture Facility (Davis, California, United States, 38.54’N, 121.87’W) during one tomato growing season in 2021. Sampling was conducted at four time points: March (pre-planting; T0; 0 weeks), June (vegetative phase, 6 weeks post-planting; T1; 9 weeks since T0), July (Flowering, 10 weeks post-planting; T2; 14 weeks since T0), and August (maturity, harvest, 16 weeks post-planting; T3; 19 weeks since T0). Three plots (64×64 m) with the Heinz 1662 tomato cultivar were sampled at each time point. In total, 78 soil samples were collected. At the pre-planting time point (March), six bulk soil samples were collected (two per plot). At each post-planting time point (June, July, and August), 24 samples were collected, comprising bulk and rhizosphere soils from two plants per plot under two treatment conditions (with and without arbuscular mycorrhizal fungi, AMF). Across post-planting time points, this resulted in sampling of three plots × two soil compartments (bulk and rhizosphere) × two plants per treatment × two treatments. Each plot was conventionally managed and left fallow during the preceding winter months, then received mineral fertilizer (urea ammonium nitrate solution; 156 kg haLJ¹ total) via drip line irrigation 3-4 times throughout the growing season. Tomato seedlings with the first two true leaves were transplanted into the field. All seedlings assigned to the AMF treatment were inoculated using a root-dip method with EndoMaxx Prime (Valent, San Ramon, CA, USA) at transplanting, following the manufacturer’s instructions. Control seedlings were dip-inoculated with autoclaved EndoMaxx Prime. Fertilization and field management were identical across treatments. No soil biocides (ie. fungicides) were applied to allow AMF establishment and soil microbiome characterization; no disease pressure was observed. At each post-planting sampling time point (June, July, August), two plants per plot were randomly selected for each treatment (with and without AMF), and their rhizosphere (soil adhering to roots after excavation) and accompanying bulk soils (30 cm from the base of the plant) were collected and processed according to the descriptions below. At the pre-planting time point (March), no plants were present and only bulk soil was sampled, two per plot, in approximately the same location (i.e., 30 cm from where plants were to be planted). Bulk soil samples were collected using a soil knife from the upper approx. 25 cm of soil (2.5 × 2.5 cm sampling area), manually homogenized, and stored at 4 °C until processing, which occurred within 24 h of collection. For rhizosphere samples, plants were excavated and roots with adhering soil were collected, homogenized, and stored at 4 °C until processing.

### Soil and plant chemistry

Bulk soil moisture was defined by measuring soil gravimetric water content. Soil physicochemical properties for bulk soil samples at all timepoints were measured by Ward Laboratories (Kearney, NE, USA). Measured variables included soil pH, buffer pH, soluble salts, organic matter, nitrate, phosphorus, potassium, sulfate, and a suite of exchangeable cations and micronutrients using standard methods (Table S1).

Soil pH and soluble salts were measured using a 1:1 mixture, by weight, of 1 part soil to 1 part distilled H_2_O (Eckert, 1988). Soil organic matter was determined as percent mass loss on ignition after heating sample in a muffle furnace to 500°C (Heiri et al., 2001). Nitrate and Ammonium were measured by colorimetry on 2M KCl extracts (Keeney & Nelson, 1982). Potassium, calcium, magnesium, and sodium were quantified using ammonium acetate extractions (Normandin et al., 1998). Zinc, iron, manganese, and copper were measured using a diethylenetriaminepentaacetic acid (DTPA) extraction (Liang & Karamanos, 1993). Phosphorus was measured using Olsen bi-carbonate solution extraction (Sims, 2000), and sulphite (sulfur trioxide) was measured using a Mehlich-3 extraction (Mehlich, 1984). Cation exchange capacity (CEC) and percent base saturation values were calculated from the sum of extracted cations.

Plant growth, root AMF colonization and fruit quality parameters were assessed at different time points throughout the growing season. Root length colonized by arbuscular mycorrhizal fungi was quantified using a gridline intersect method (Giovannetti & Mosse, 1980) following root staining (Vierheilig et al., 1998). Briefly, roots were rinsed in PBS after rhizosphere soil collection, subsampled (0.1 g) and stored at 4°C. Root subsamples were then cut into approximately 2-centimeter segments and stained using a method modified from Vierheilig et al. (1998). Root segments were cleared in a 10% KOH solution at 90°C for 40 minutes and stained in a 5% ink and vinegar solution for 5 minutes at 90°C. Roots were then bleached in 10% sodium hypochlorite solution for 5 minutes. Roots were stained in 5% ink-vinegar solution and stored in 10% water solution prior to quantification. AMF colonization was visually quantified using the gridline intersect method (Giovannetti and Mosse, 1980) on a dissecting scope at x 80 magnification. A minimum of 100 root segments were counted for each sample.

Vine biomass and fruit yield estimates were obtained through destructive sampling. Dry vine biomass (kg/ha) was determined by harvesting vines from subsampled plants, drying at 60°C for four weeks, and extrapolating to plot-level area. Red, green, and total fruit hand harvest estimates (MT/ha) were calculated by collecting fruit from six randomly selected plants per plot, and extrapolating yields to the entire row based on the proportion of sampled distance to total row length. Total fruit yield was calculated as the sum of red and green fruit. A subsample of red fruits was collected from each subplot and taken to a Processing Tomato Advisory Board (http://www.ptab.org) grading station to measure total soluble solids, fruit color and fruit pH. Total soluble solids (°Brix) were measured using a refractometer on collected fruit samples. Fruit pH was determined on blended fruit samples as described above. Fruit color was quantified as Hunter hue angle using a Konica Minolta model CR-410 colorimeter on whole tomatoes blended under high vacuum to remove air bubbles.

Plant tissue composed of three leaflets per plants mixed to form a composite sample representative of each plot were analyzed for macro-and micronutrient concentrations at Pennsylvania State University for nutrient analyses (State College, PA). Total nitrogen content (%) was determined by thermal conductivity using combustion analysis with an Elementar Vario Max N/C Analyzer, where dried plant material ignited in an induction furnace at approximately 900°C in a helium and oxygen environment within quartz tube (Horneck & Miller, 1998). Phosphorus, potassium, calcium, magnesium, and sulfur (%) along with manganese, iron, copper, boron, aluminum, zinc, and sodium (mg/kg) were determined following acid digestion (Huang & Schulte, 1985). Dried, ground plant material was mixed with an internal standard and pre-digestant, heated to 60°C in an aluminum block, digested with acid at 120°C, then diluted, filtered, and analyzed using ICP emission spectroscopy.

### Virome DNA extraction, library construction, and shotgun sequencing

Samples were processed within 24 hours after collection for viromics and kept at 4 °C until processing. For each bulk soil sample, 10 grams of soil were suspended in 30 mL of protein-supplemented phosphate-buffered saline solution (PPBS: 2% bovine serum albumin, 10% phosphate-buffered saline, 1% potassium citrate, and 150 mM MgSO_4_) and then briefly vortexed and placed on an orbital shaker (30 min, 400 rpm, 4 °C). For each rhizosphere sample, the roots were vigorously shaken to remove loose soil particles, and only the adhered portion was analyzed. The roots were suspended and shaken in the same PPBS buffer as for bulk soils, with no explicit differentiation between the rhizosphere and endosphere (i.e., the cut ends of roots were also exposed to the buffer). From this point, bulk soil and rhizosphere virome processing was the same. The shaken soil-buffer solution was centrifuged (10 min, 3,095 x g, 4 °C) and the resulting supernatant was then centrifuged twice (8 min, 10,000 x g, 4 °C) to remove residual soil particles. The centrifuged supernatants were then filtered through a 0.22 μm polyethersulfone membrane filter to remove most cells. The filtrate was then ultracentrifuged (2 hrs 25 min, 32,000 x rpm, 4 °C) to pellet the virions, using an Optima LE-80K ultracentrifuge with a 50.2 Ti rotor (Beckman-Coulter Life Sciences). Supernatant was discarded, and pellets were resuspended in 100 μl of ultrapure water and treated with 10 units of RQ1 RNase-free DNase and 10 μl of 10× DNase buffer (Promega Corp., Madison, WI, USA) to remove free DNA. Samples were incubated at 37 °C for 30 min, and the reaction was stopped by adding 10 μl of the DNase stop solution (Promega Corp., Madison, WI, USA). DNA was then extracted from the viral fraction, using the DNeasy PowerSoil Pro kit (Qiagen, Hilden, Germany), following the manufacturer’s instructions, with an added step of a 10-minute incubation at 65 °C before the bead-beating step. Libraries were constructed using the DNA Hyper Prep library kit (Kapa Biosystems-Roche, Basel, Switzerland). Paired-end 150 bp sequencing was done using the NovaSeq S4 platform (Illumina), to an approximate depth of 10 Gbp per virome.

### Total DNA extraction, amplicon library construction, and sequencing

For each bulk soil sample, total DNA was extracted from 0.25 g of soil with the DNeasy PowerSoil Pro kit (Qiagen, Hilden, Germany), following the manufacturer’s instructions, with an added step of a 10-minute incubation at 65 °C prior to the bead-beating step. For each rhizosphere sample, 0.25 g of soil was brushed off the roots into the extraction tube, followed by the same extraction protocol. Construction of the amplicon libraries followed a previously described dual-indexing strategy (Caporaso et al., 2010; Edwards et al., 2018). To target the V4 region of the 16S rRNA gene, universal primers 515F and 806R were used, using the following PCR protocol: an initial denaturation step at 98 °C for 2 min, followed by 30 cycles of 98 °C for 20 s, 50 °C for 30 s and 72 °C for 45 s, and a final extension step at 72 °C for 10 min. To amplify the ITS1 region, we used the universal primers ITS1-F and ITS2 (Agler et al., 2016; Gardes & Bruns, 1993; White et al., 1990) and the following PCR program: an initial denaturation step at 95 °C for 2 min, followed by 35 cycles of 95 °C for 20 s, 50 °C for 30 s, and 72 °C for 50 s, followed by a final extension at 72 °C for 10 min. All PCR reactions were performed using the Platinum Hot Start PCR Master Mix (Invitrogen). Libraries were cleaned using AmpureXP magnetic beads (Beckman-Coulter Life Sciences), quantified (Qubit 4 fluorometer, ThermoFisher Scientific), and pooled in equimolar concentrations. Paired-end sequencing (250 bp) was performed on the MiSeq platform (Illumina), using a standard flow cell per library (one for 16S rRNA gene amplicons and one for ITS amplicons).

### RNA extraction, library construction, and sequencing

Plants were excavated from the soil and vigorously shaken. Roots with adhering soil were collected as rhizosphere samples. Each rhizosphere sample was divided into three portions: one portion was kept at 4°C (on ice) for DNA extraction (as described above), another was flash-frozen in liquid nitrogen for RNA extraction, and a third portion was reserved for arbuscular mycorrhizal fungal (AMF) colonization assessment via root staining. Rhizosphere samples designated for RNA extraction were immediately flash-frozen in liquid nitrogen in the field and stored at-80°C until further processing. Total RNA was extracted using the RNeasy PowerSoil Pro Kit (Qiagen, Hilden, Germany), following the manufacturer’s instructions. RNA was submitted to Genewiz (San Francisco, CA, USA) for ribodepletion, cDNA preparation, library construction via the NEBNext Ultra II RNA library prep kit (New England Biolabs, Ipswitch, MA, USA), and sequencing. Paired-end sequencing (150 bp) was done using the NovaSeq 6000 platform (Illumina) to an approximate sequencing depth of 10 Gbp per sample.

### DNA virome bioinformatic processing

Virome sequencing reads were trimmed using Trimmomatic v0.39 (Bolger et al., 2014), removing Illumina adapters and quality trimming reads, using paired-end trimming, a sliding window size of 3:40, and a minimum read length of 50 bp. PhiX sequences were removed using bbduk from the BBMap v38.72 package (Bushnell, 2014), using k=31 and hdist=1, and host plant reads were removed using Bowtie2 v2.4.2 (Langmead & Salzberg, 2012), using the sensitive setting, mapping against the genome of tomato (*S. lycopersicum*),

GenBank accession number GCA_012431665. The remaining reads were assembled into contigs, using MEGAHIT 1.0.6 (D. Li et al., 2015), with settings k-min 27, minimum contig length of 10 kbp, and presets meta-large. Resulting contigs were renamed using the rename function from BBMap, and viral contigs were predicted using VIBRANT v1.2.0 (Kieft et al., 2020), in virome mode. All predicted viral contigs were dereplicated into vOTUs, using dRep v3.2.0 (Olm et al., 2017), at 95% average nucleotide identity (ANI) with a minimum alignment of 85%, using the ANImf algorithm. Reads were mapped to the vOTUs and to the PIGEONv2.0 database (ter Horst, Fudyma, Sones, et al., 2023) using Bowtie2 v2.4.2 (Langmead & Salzberg, 2012), using sensitive mode. vOTUs from the PIGEONv2.0 database that were recovered in this dataset were clustered with the vOTUs assembled from this dataset using dRep, using the same settings as above. Reads were then again mapped to this non-redundant dataset of vOTUs, using Bowtie2, and the resulting samfiles were converted to bamfiles using SAMtools v1.15.1 (H. Li et al., 2009). A coverage table was created with CoverM v0.6.1 (Aroney et al., 2025) using CoverM contig with the mean coverage and a minimum covered fraction (breadth) of 75%. The resulting coverage table was used for statistical analysis, unless otherwise noted. All scripts for bioinformatic processing are available on Github (https://github.com/AnneliektH/TomatoRhizo).

### Amplicon sequence bioinformatic processing

Paired-end reads were assembled into single sequences using PANDAseq v2.9 (Masella et al., 2012). Chimeric sequences were removed, reads were dereplicated, and denoising and read merging were performed using DADA2 v1.12.1 with default settings, following the standard DADA2 workflow ((Callahan et al., 2016); https://benjjneb.github.io/dada2/tutorial.html). Taxonomy was assigned using the RDP classifier implementation in DADA2 (Wang et al., 2007) with the SILVA database v132 (Quast et al., 2013) for 16S rRNA gene sequences, and with the UNITE database v2021-05-10 for ITS sequences (Abarenkov et al., 2020). Amplicon sequence variant (ASV) tables with raw read counts per sample were generated using the ‘makeSequenceTablè function in DADA2. Sequences matching mitochondria or chloroplasts (0.11% of total 16S rRNA gene reads) were removed from the 16S rRNA gene dataset using phyloseq v.1.52.0 (McMurdie & Holmes, 2013). ASV tables were converted to relative abundances prior to downstream analyses.

### Metatranscriptome bioinformatic processing for both RNA viral community and community transcript analyses

Paired-end metatranscriptomic reads were first trimmed, and PhiX and the host plant (tomato) were removed, following the same procedures used for DNA viromes above. The remaining reads were assembled into contigs using MEGAHIT v1.0.6 (D. Li et al., 2015), with settings k-min 27, minimum contig length of 200 bp, and presets meta-large. Predicted proteins were generated from assembled contigs using Prodigal v2.6.3 (Hyatt et al., 2010), using standard settings. For RNA viral analyses, HMMER v3.3.2 (Eddy, 2011) was used to recover RNA-dependent RNA polymerase (RdRp) sequences with a p-value threshold of 1e-05, as previously described (Starr et al., 2019; ter Horst, Fudyma, Bak, et al., 2023). Predicted proteins were further annotated using DIAMOND v0.9.22.123 (Buchfink et al., 2015) against the NCBI nr prokaryotic database (v203), using a p-value threshold of 1e-06. Contigs containing prokaryotic genes were flagged and reads mapping to these contigs were removed to reduce prokaryotic RNA signal. The remaining reads were re-assembled using MEGAHIT, and this process was iterated three times. The final set of contigs containing predicted RdRp sequences was clustered using dRep v3.2.0 (Olm et al., 2017) at 95% ANI with a minimum alignment of 85%, using the ANImf algorithm. Metatranscriptomic reads were mapped back to these sequences and to RefSeq v203 RNA viral sequences (n=4,472) using Bowtie2, and a coverage table was generated using CoverM v0.6.1 (Aroney et al., 2025) with the same parameters as described for DNA viromes.

For community transcriptome analyses, reads remaining after host (tomato) removal were further processed to reduce rRNA contamination. rRNA reads were removed using SortMeRNA v4.3.7 (Kopylova et al., 2012) with default rRNA reference databases (v4.3.4), and unaligned reads were retained for downstream analyses. Because SortMeRNA outputs interleaved reads, BBTools repair.sh (Bushnell, 2014) was used to restore separate forward and reverse read files. Non-rRNA reads were then mapped to a reference transcriptome, generated by co-assembly of host filtered reads using MEGAHIT, as described above. Salmon v1.10.3 index (Patro et al., 2017) of contigs was built in quasi-mapping mode, and transcript abundance was quantified per sample using Salmon with automatic library type detection and default parameters. Salmon outputs transcript-level read estimates that account for sequence composition and multi-mapping reads (Patro et al., 2017). Quantification outputs from all samples were merged using Salmon quantmerge to generate a matrix of raw read counts for downstream analyses. For functional annotation, open reading frames (ORFs) were predicted from contigs using TransDecoder v5.7.1 (Haas, 2024), and long ORFs were translated into protein sequences. Proteins were annotated using eggNOG-mapper v2.1.13 (Cantalapiedra et al., 2021). Contig-level functional annotations were linked to transcript abundance for downstream analyses. Community metatranscriptome data processing scripts are available on Github (https://github.com/ljiraska/Tomato_rhizo_AMF).

### Statistical analysis

All statistical analyses were performed using R v4.5.2 (R Core Team, 2025), using the mean coverage vOTU abundance table or the other coverage tables prepared as described above (for RdRps, transcript table, 16S rRNA gene ASVs, and ITS ASVs, respectively), and ggplot2 v package (Wickham et al., 2024), unless otherwise noted. Phyloseq objects were prepared with abundance tables and metadata using R package phyloseq v.1.52.0 (McMurdie & Holmes, 2013). ASV tables were CSS normalized and all tables were Hellinger transformed with transform function from microbiome v1.30.0 (Lahti & Shetty, 2022) package. Statistical significance was defined as p < 0.05 for single comparisons and adjusted p < 0.05 (Benjamini-Hochberg correction) for multiple comparisons unless otherwise noted.

Alpha-diversity metrics (observed richness) were calculated using microbiome v1.30.0 (Lahti & Shetty, 2022) using function alpha. Statistical significance between bulk and rhizosphere for observed richness, and DNA concentration was assessed using rstatix v0.7.2 (Kassambara, 2025) using wilcox_test function for a Wilcoxon rank-sum test and for plant related variables using t_test function for t-test.

Principal coordinate analysis (PCoA) with Bray-Curtis dissimilarity matrices were calculated for viral, bacterial and fungal communities using the ordinate function from the phyloseq package v1.52.0 (McMurdie & Holmes, 2013). Canonical analysis of principal coordinates (CAP) was conducted to assess the effect of plot on community structure using Bray-Curtis distances with the formula ∼Plot, also with ordinate function. Soil chemistry and plant chemistry data were each standardized (mean-centered and scaled to unit variance) prior to PCA using the scale function in base R. Moisture content was excluded from soil chemistry analysis for PCA and PCA was performed with prcomp function in base R.

Permutational multivariate analysis of variance (PERMANOVA) was conducted using the adonis2 function from the vegan package v2.7.2 (Oksanen et al., 2022) to test the effects of sample type (bulk vs. rhizosphere), treatment, timepoint, and plot on community composition based on Bray-Curtis dissimilarity matrices and using scaled soil chemistry and plant data with Euclidean distance matrices. PERMANOVA was performed with 999 permutations, and effects were evaluated sequentially by terms. Where appropriate, two-and three-way interactions were also tested. Mantel tests were performed to assess correlations between community (using Bray-Curtis) and soil chemistry (Euclidean) dissimilarity matrices of different organisms using the mantel function from the vegan package with 999 permutations and Pearson correlation.

Linear regressions and Pearson’s correlation tests were performed to assess relationships between PCoA axis 1 scores and environmental variables (e.g., soil moisture) using the cor.test function in base R.

Variation partitioning analysis was conducted following the approach described by Legendre and Legendre (2012) to decompose the variation in community composition explained by soil moisture, selected soil chemistry variables, location of sampling, and temporal factors. Forward selection of soil chemistry variables was performed using the forward.sel function from the adespatial v0.3.28 package, with the adjusted R² threshold set to the global adjusted R² of the full soil chemistry model and 999 permutations. When forward selection identified significant variables, only these selected variables were retained for soil chemistry in the subsequent variation partitioning. Variation partitioning was performed using the varpart function from the vegan package. For temporal effects, variation partitioning was performed separately using categorical timepoint (four discrete sampling dates) and linear time (weeks since first sampling) to compare discrete versus continuous temporal models.

Heatmaps of scaled bulk soil chemistry data were generated using the pheatmap function from the pheatmap package v1.0.13 (Kolde, 2025) with hierarchical clustering of samples based on Euclidean distance (default settings).

Differential gene expression analysis was performed using DESeq2 v1.48.2 (Love et al., 2023), following the methods described in the tutorial of the package vignette (https://bioconductor.org/packages/devel/bioc/vignettes/DESeq2/inst/doc/DESeq2.html#principal-component-plot-of-the-samples). A DESeqDataSet object was constructed with a design formula accounting for site, timepoint, and treatment effects (∼ Site + Timepoint + Treatment). Differential expression testing was conducted using the default Wald test, comparing AMF treatment versus control samples. Log2 fold changes were shrunk using the apeglm method to reduce noise from low-count genes. Transcripts with adjusted p-value < 0.05 (Benjamini-Hochberg correction) and absolute log2 fold change > 1 were considered differentially expressed genes. Functional annotation of transcripts was obtained from eggNOG-mapper (as described above), with isoform suffixes removed and the best annotation per transcript retained based on lowest e-value. For community gene expression data, principal component analysis (PCA) was performed on variance-stabilized transformed count data using the plotPCA function from DESeq2 v1.48.2 (Love et al., 2023). The analysis included timepoint, treatment, and site as grouping variables.

To evaluate desiccation stress in rhizosphere microbial communities, we used 22 previously identified desiccation-upregulated genes in soil bacterium *Curtobacterium* (Scales et al., 2023). We filtered metatranscriptomic data to genes including those involved in osmolyte synthesis (dhaK, dhaM, treZ, treY, otsB), DNA repair and recombination (recA, recN, uvrA, dps), SOS response (lexA, recX), oxidative stress response (katA, ahpF), efflux pumps (macB, arsB, cnrH), sporulation and stress sigma factors (gdh, sigK, rpoE), and other desiccation-related functions (tgnD, gabD, fadA) (Scales et al., 2023). Relative abundance of desiccation genes was calculated as the sum of desiccation gene transcript abundance divided by total transcript abundance per sample. Temporal trends were analyzed using linear mixed-effects models with lme4 v1.1.37 package (Bates et al., 2022) with time (weeks) and treatment, as fixed effects and plot location as a random effect. P-values were obtained using the lmerTest v3.1.3 package (Kuznetsova et al., 2026). Treatment-specific temporal slopes were calculated using the emmeans v1.11.2.8 package (Lenth et al., 2025).

All maps were made using the R packages rnaturalearth v1.2.0 (Massicotte et al., 2026), rnaturalearthdata v1.0.0 (South et al., 2024) and sf v1.0.21 (Pebesma et al., 2026), and all other plots were created using the R package ggplot2 v4.0.0 (Wickham, Chang, et al., 2026), and eulerr v7.0.4 (Larsson et al., 2025). Plots were composed together with R package patchwork v1.3.2 (Pedersen, 2025) and Affinity Designer 2 was used to make modification to improve readability of figures. For data manipulation we used R packages tidyverse v2.0.0 (Wickham & RStudio, 2023), dplyr v1.1.4 (Wickham, François, et al., 2026) and stringr v1.5.2 (Wickham et al., 2025). All data processing and statistical analysis scripts are available on Github (https://github.com/ljiraska/Tomato_rhizo_AMF).

## Data availability

Raw sequencing reads for viromes, metatranscriptomes, 16S rRNA gene, ITS amplicon libraries, are available on NCBI under BioProject PRJNA937255. Final set of vOTU sequences and RdRps are available on Dryad (https://datadryad.org/), using the following https://doi.org/10.5061/dryad.sf7m0cgnb. vOTU sequences are also available on Dryad within the PIGEONv2.0 database (https://doi.org/10.25338/B8C934).

## Acknowledgements

This study was primarily supported by USDA NIFA AFRI award # 2021-144 67013-34815-0 (grant to JBE). Additional support was provided by USDA NIFA Hatch # CA-D145 PPA-2464-H. We thank Simon Roux and Rex Malmstrom (DOE LBNL JGI) for their exploratory density fractionation and sequencing of viromic DNA from two Russell Ranch samples from 2018, which became part of our PIGEONv2.0 database and which contributed to vOTU recovery via read mapping in this study. We thank Jess Sorensen and Sarah Geonczy for help with sampling.

## References

Abarenkov, K., Zirk, A., Piirmann, T., Pöhönen, R., Ivanov, F., Nilsson, R. H., & Kõljalg, U. (2020). UNITE QIIME release for eukaryotes 2 (Version 8.2) [Computer software]. 10.15156/BIO/786388

Adriaenssens, E. M., Kramer, R., Van Goethem, M. W., Makhalanyane, T. P., Hogg, I., & Cowan, D. A. (2017). Environmental drivers of viral community composition in Antarctic soils identified by viromics. Microbiome, 5(1), 83. 10.1186/s40168-017-0301-7

Agler, M. T., Ruhe, J., Kroll, S., Morhenn, C., Kim, S.-T., Weigel, D., & Kemen, E. M. (2016). Microbial Hub Taxa Link Host and Abiotic Factors to Plant Microbiome Variation. PLoS Biology, 14(1), e1002352. 10.1371/journal.pbio.1002352

Aroney, S. T. N., Newell, R. J. P., Nissen, J. N., Camargo, A. P., Tyson, G. W., & Woodcroft, B. J. (2025). CoverM: Read alignment statistics for metagenomics. Bioinformatics, 41(4), btaf147. 10.1093/bioinformatics/btaf147

Bakker, M. G., Chaparro, J. M., Manter, D. K., & Vivanco, J. M. (2015). Impacts of bulk soil microbial community structure on rhizosphere microbiomes of Zea mays. Plant and Soil, 392(1), 115–126. 10.1007/s11104-015-2446-0

Bakker, P. A. H. M., Berendsen, R. L., Doornbos, R. F., Wintermans, P. C. A., & Pieterse, C. M. J. (2013). The rhizosphere revisited: Root microbiomics. Frontiers in Plant Science, 4. 10.3389/fpls.2013.00165

Bakker, P. A. H. M., Pieterse, C. M. J., Jonge, R. de, & Berendsen, R. L. (2018). The Soil-Borne Legacy. Cell, 172(6), 1178–1180. 10.1016/j.cell.2018.02.024

Balliu, A., Sallaku, G., & Rewald, B. (2015). AMF Inoculation Enhances Growth and Improves the Nutrient Uptake Rates of Transplanted, Salt-Stressed Tomato Seedlings. Sustainability, 7(12), Article 12. 10.3390/su71215799

Banerjee, S., & van der Heijden, M. G. A. (2023). Soil microbiomes and one health. Nature Reviews Microbiology, 21(1), 6–20. 10.1038/s41579-022-00779-w

Bates, D., Maechler, M., Bolker [aut, B., cre, Walker, S., Christensen, R. H. B., Singmann, H., Dai, B., Scheipl, F., Grothendieck, G., Green, P., Fox, J., Bauer, A., & simulate.formula), P. N. K. (shared copyright on. (2022). lme4: Linear Mixed-Effects Models using “Eigen” and S4 (Version 1.1-30) [Computer software]. https://CRAN.R-project.org/package=lme4

Benard, P., Zarebanadkouki, M., Brax, M., Kaltenbach, R., Jerjen, I., Marone, F., Couradeau, E., Felde, V. J. M. N. L., Kaestner, A., & Carminati, A. (2019). Microhydrological Niches in Soils: How Mucilage and EPS Alter the Biophysical Properties of the Rhizosphere and Other Biological Hotspots. Vadose Zone Journal, 18(1), 180211. 10.2136/vzj2018.12.0211

Bi, L., Yu, D.-T., Du, S., Zhang, L.-M., Zhang, L.-Y., Wu, C.-F., Xiong, C., Han, L.-L., & He, J.-Z. (2021). Diversity and potential biogeochemical impacts of viruses in bulk and rhizosphere soils. Environmental Microbiology, 23(2), 588–599. 10.1111/1462-2920.15010

Bin Jang, H., Bolduc, B., Zablocki, O., Kuhn, J. H., Roux, S., Adriaenssens, E. M., Brister, J. R., Kropinski, A. M., Krupovic, M., Lavigne, R., Turner, D., & Sullivan, M. B. (2019). Taxonomic assignment of uncultivated prokaryotic virus genomes is enabled by gene-sharing networks. Nature Biotechnology, 37(6), 632–639. 10.1038/s41587-019-0100-8

Bolger, A. M., Lohse, M., & Usadel, B. (2014). Trimmomatic: A flexible trimmer for Illumina sequence data. Bioinformatics, 30(15), 2114–2120. 10.1093/bioinformatics/btu170

Brockett, B. F. T., Prescott, C. E., & Grayston, S. J. (2012). Soil moisture is the major factor influencing microbial community structure and enzyme activities across seven biogeoclimatic zones in western Canada. Soil Biology and Biochemistry, 44(1), 9–20. 10.1016/j.soilbio.2011.09.003

Buchfink, B., Xie, C., & Huson, D. H. (2015). Fast and sensitive protein alignment using DIAMOND. Nature Methods, 12(1), 59–60. 10.1038/nmeth.3176

Bulgarelli, D., Schlaeppi, K., Spaepen, S., Themaat, E. V. L. van, & Schulze-Lefert, P. (2013). Structure and Functions of the Bacterial Microbiota of Plants. Annual Review of Plant Biology, 64(Volume 64, 2013), 807–838. 10.1146/annurev-arplant-050312-120106

Bushnell, B. (2014). BBMap: A Fast, Accurate, Splice-Aware Aligner (LBNL-7065E). Lawrence Berkeley National Lab. (LBNL), Berkeley, CA (United States). https://www.osti.gov/biblio/1241166

Callahan, B. J., McMurdie, P. J., Rosen, M. J., Han, A. W., Johnson, A. J. A., & Holmes, S. P. (2016). DADA2: High-resolution sample inference from Illumina amplicon data. Nature Methods, 13(7), 581–583. 10.1038/nmeth.3869

Cantalapiedra, C. P., Hernández-Plaza, A., Letunic, I., Bork, P., & Huerta-Cepas, J. (2021). eggNOG-mapper v2: Functional Annotation, Orthology Assignments, and Domain Prediction at the Metagenomic Scale. Molecular Biology and Evolution, 38(12), 5825–5829. 10.1093/molbev/msab293

Cao, M.-M., Liu, S.-Y., Bi, L., Chen, S.-J., Wu, H.-Y., Ge, Y., Han, B., Zhang, L.-M., He, J.-Z., & Han, L.-L. (2022). Distribution Characteristics of Soil Viruses Under Different Precipitation Gradients on the Qinghai-Tibet Plateau. Frontiers in Microbiology, 13. 10.3389/fmicb.2022.848305

Caporaso, J. G., Kuczynski, J., Stombaugh, J., Bittinger, K., Bushman, F. D., Costello, E. K., Fierer, N., Peña, A. G., Goodrich, J. K., Gordon, J. I., Huttley, G. A., Kelley, S. T., Knights, D., Koenig, J. E., Ley, R. E., Lozupone, C. A., McDonald, D., Muegge, B. D., Pirrung, M.,…Knight, R. (2010). QIIME allows analysis of high-throughput community sequencing data. Nature Methods, 7(5), 335–336. 10.1038/nmeth.f.303

Carminati, A., Zarebanadkouki, M., Kroener, E., Ahmed, M. A., & Holz, M. (2016). Biophysical rhizosphere processes affecting root water uptake. Annals of Botany, 118(4), 561–571. 10.1093/aob/mcw113

Chaparro, J. M., Sheflin, A. M., Manter, D. K., & Vivanco, J. M. (2012). Manipulating the soil microbiome to increase soil health and plant fertility. Biology and Fertility of Soils, 48(5), 489–499. 10.1007/s00374-012-0691-4

Chariou, P. L., Dogan, A. B., Welsh, A. G., Saidel, G. M., Baskaran, H., & Steinmetz, N. F. (2019). Soil mobility of synthetic and virus-based model nanopesticides. Nature Nanotechnology, 14(7), 712–718. 10.1038/s41565-019-0453-7

Chen, Y., Sun, C., Yan, Y., Jiang, D., Huangfu, S., & Tian, L. (2025). Impact of arbuscular mycorrhizal fungi on maize rhizosphere microbiome stability under moderate drought conditions. Microbiological Research, 290, 127957. 10.1016/j.micres.2024.127957

Das, P. P., Singh, K. R., Nagpure, G., Mansoori, A., Singh, R. P., Ghazi, I. A., Kumar, A., & Singh, J. (2022). Plant-soil-microbes: A tripartite interaction for nutrient acquisition and better plant growth for sustainable agricultural practices. Environmental Research, 214, 113821. 10.1016/j.envres.2022.113821

DiPietro, A. G., Bryant, S. A., Zanger, M. M., & Williamson, K. E. (2023). Understanding Viral Impacts in Soil Microbial Ecology Through the Persistence and Decay of Infectious Bacteriophages. Current Microbiology, 80(9), 276. 10.1007/s00284-023-03386-x

Durham, D. M., Sieradzki, E. T., ter Horst, A. M., Santos-Medellín, C., Bess, C. W. A., Geonczy, S. E., & Emerson, J. B. (2022). Substantial differences in soil viral community composition within and among four Northern California habitats. ISME Communications, 2(1), 100. 10.1038/s43705-022-00171-y

E. Evans, S., D. Allison, S., & V. Hawkes, C. (2022). Microbes, memory and moisture: Predicting microbial moisture responses and their impact on carbon cycling. Functional Ecology, 36(6), 1430–1441. 10.1111/1365-2435.14034

Eckert, D. J. (1988). Recommended pH and lime requirement tests. Recommended Chemical Soil Test Procedures for the North Central Region. North Central Regional Publication, 221, 6–8.

Eddy, S. R. (2011). Accelerated Profile HMM Searches. PLOS Computational Biology, 7(10), e1002195. 10.1371/journal.pcbi.1002195

Edwards, J., Johnson, C., Santos-Medellín, C., Lurie, E., Podishetty, N. K., Bhatnagar, S., Eisen, J. A., & Sundaresan, V. (2015). Structure, variation, and assembly of the root-associated microbiomes of rice. Proceedings of the National Academy of Sciences, 112(8), E911–E920. 10.1073/pnas.1414592112

Edwards, J., Santos-Medellín, C., & Sundaresan, V. (2018). Extraction and 16S rRNA Sequence Analysis of Microbiomes Associated with Rice Roots. Bio-Protocol, 8(12), e2884. 10.21769/BioProtoc.2884

Emerson, J. B. (2019). Soil Viruses: A New Hope. mSystems, 4(3), 10.1128/msystems.00120-19

Emerson, J. B., Roux, S., Brum, J. R., Bolduc, B., Woodcroft, B. J., Jang, H. B., Singleton, C. M., Solden, L. M., Naas, A. E., Boyd, J. A., Hodgkins, S. B., Wilson, R. M., Trubl, G., Li, C., Frolking, S., Pope, P. B., Wrighton, K. C., Crill, P. M., Chanton, J. P.,…Sullivan, M. B. (2018). Host-linked soil viral ecology along a permafrost thaw gradient. Nature Microbiology, 3(8), 870–880. 10.1038/s41564-018-0190-y

Fierer, N. (2017). Embracing the unknown: Disentangling the complexities of the soil microbiome. Nature Reviews Microbiology, 15(10), 579–590. 10.1038/nrmicro.2017.87

Fierer, N., Breitbart, M., Nulton, J., Salamon, P., Lozupone, C., Jones, R., Robeson, M., Edwards, R. A., Felts, B., Rayhawk, S., Knight, R., Rohwer, F., & Jackson, R. B. (2007). Metagenomic and Small-Subunit rRNA Analyses Reveal the Genetic Diversity of Bacteria, Archaea, Fungi, and Viruses in Soil. Applied and Environmental Microbiology, 73(21), 7059–7066. 10.1128/AEM.00358-07

Fudyma, J. D., Penev, P., Estera-Molina, K., Hoff, J., Blazewicz, S. J., Pett-Ridge, J., & Emerson, J. B. (2025). Spatial and depth structuring predominate over temporal variation in Mediterranean grassland soil viral communities (p. 2025.12.30.696869). bioRxiv. 10.64898/2025.12.30.696869

Fudyma, J. D., ter Horst, A. M., Santos-Medellín, C., Sorensen, J. W., Gogul, G. G., Hillary, L. S., Geonczy, S. E., Pett-Ridge, J., & Emerson, J. B. (2024). Exploring viral particle, soil, and extraction buffer physicochemical characteristics and their impacts on extractable viral communities. Soil Biology and Biochemistry, 194, 109419. 10.1016/j.soilbio.2024.109419

Gardes, M., & Bruns, T. D. (1993). ITS primers with enhanced specificity for basidiomycetes—Application to the identification of mycorrhizae and rusts. Molecular Ecology, 2(2), 113–118. 10.1111/j.1365-294X.1993.tb00005.x

Geonczy, S. E., Hillary, L. S., Santos-Medellín, C., Sorensen, J. W., & Emerson, J. B. (2025). Patchy burn severity explains heterogeneous soil viral and prokaryotic responses to fire in a mixed conifer forest. mSystems, 10(6), e01749–24. 10.1128/msystems.01749-24

Geonczy, S. E., ter Horst, A. M., & Emerson, J. B. (2025). Soil viral communities shifted significantly after wildfire in chaparral and woodland habitats. ISME Communications, 5(1), ycaf073. 10.1093/ismeco/ycaf073

Gerba, C. P. (1984). Applied and Theoretical Aspects of Virus Adsorption to Surfaces. In Advances in Applied Microbiology (Vol. 30, pp. 133–168). Academic Press. 10.1016/S0065-2164(08)70054-6

Giovannetti, M., & Mosse, B. (1980). An evaluation of techniques for measuring vesicular arbuscular mycorrhizal infection in roots. New Phytologist, 489–500.

Gosling, P., Hodge, A., Goodlass, G., & Bending, G. D. (2006). Arbuscular mycorrhizal fungi and organic farming. Agriculture, Ecosystems & Environment, 113(1), 17–35. 10.1016/j.agee.2005.09.009

Guzman, A., Montes, M., Hutchins, L., DeLaCerda, G., Yang, P., Kakouridis, A., Dahlquist-Willard, R. M., Firestone, M. K., Bowles, T., & Kremen, C. (2021). Crop diversity enriches arbuscular mycorrhizal fungal communities in an intensive agricultural landscape. New Phytologist, 231(1), 447–459. 10.1111/nph.17306

Haas, B. J. (2024). TransDecoder (Version 5.7.1) [Computer software]. https://github.com/TransDecoder/TransDecoder

Hajiboland, R., Aliasgharzadeh, N., Laiegh, S. F., & Poschenrieder, C. (2010). Colonization with arbuscular mycorrhizal fungi improves salinity tolerance of tomato (Solanum lycopersicum L.) plants. Plant and Soil, 331(1), 313–327. 10.1007/s11104-009-0255-z

Heiri, O., Lotter, A. F., & Lemcke, G. (2001). Loss on ignition as a method for estimating organic and carbonate content in sediments: Reproducibility and comparability of results. Journal of Paleolimnology, 25(1), 101–110.

Hillary, L. S., Adriaenssens, E. M., Jones, D. L., & McDonald, J. E. (2022). RNA-viromics reveals diverse communities of soil RNA viruses with the potential to affect grassland ecosystems across multiple trophic levels. ISME Communications, 2, 34. 10.1038/s43705-022-00110-x

Hinsinger, P., Gobran, G. R., Gregory, P. J., & Wenzel, W. W. (2005). Rhizosphere geometry and heterogeneity arising from root-mediated physical and chemical processes. New Phytologist, 168(2), 293–303. 10.1111/j.1469-8137.2005.01512.x

Horneck, D. A., & Miller, R. O. (1998). Determination of total nitrogen in plant tissue. Handbook of Reference Methods for Plant Analysis, 75–83.

Huang, C.-Y. L., & Schulte, E. E. (1985). Digestion of plant tissue for analysis by ICP emission spectroscopy. Communications in Soil Science and Plant Analysis, 16(9), 943–958.

Hyatt, D., Chen, G.-L., LoCascio, P. F., Land, M. L., Larimer, F. W., & Hauser, L. J. (2010). Prodigal: Prokaryotic gene recognition and translation initiation site identification. BMC Bioinformatics, 11(1), 119. 10.1186/1471-2105-11-119

Kassambara, A. (2025). rstatix: Pipe-Friendly Framework for Basic Statistical Tests (Version 0.7.3) [Computer software]. https://cran.r-project.org/web/packages/rstatix/index.html

Keeney, D. R., & Nelson, D. W. (1982). Nitrogen—Inorganic forms. Methods of Soil Analysis: Part 2 Chemical and Microbiological Properties, 9, 643–698.

Kieft, K., Zhou, Z., & Anantharaman, K. (2020). VIBRANT: Automated recovery, annotation and curation of microbial viruses, and evaluation of viral community function from genomic sequences. Microbiome, 8(1), 90. 10.1186/s40168-020-00867-0

Kimura, M., Jia, Z.-J., Nakayama, N., & Asakawa, S. (2008). Ecology of viruses in soils: Past, present and future perspectives. Soil Science and Plant Nutrition, 54(1), 1–32. 10.1111/j.1747-0765.2007.00197.x

Kolde, R. (2025). pheatmap: Pretty Heatmaps (Version 1.0.13) [Computer software]. https://cran.r-project.org/web/packages/pheatmap/index.html

Kopylova, E., Noé, L., & Touzet, H. (2012). SortMeRNA: Fast and accurate filtering of ribosomal RNAs in metatranscriptomic data. Bioinformatics, 28(24), 3211–3217. 10.1093/bioinformatics/bts611

Kuznetsova, A., Brockhoff, P. B., Christensen, R. H. B., & Jensen, S. P. (2026). lmerTest: Tests in Linear Mixed Effects Models (Version 3.2-0) [Computer software]. https://cran.r-project.org/web/packages/lmerTest/index.html

Kuzyakov, Y., & Mason-Jones, K. (2018). Viruses in soil: Nano-scale undead drivers of microbial life, biogeochemical turnover and ecosystem functions. Soil Biology and Biochemistry, 127, 305–317. 10.1016/j.soilbio.2018.09.032

Lahti, L., & Shetty, S. (2022). *microbiome: Microbiome Analytics* (Version 1.18.0) [Computer software]. Bioconductor version: Release (3.15). 10.18129/B9.bioc.microbiome

Langmead, B., & Salzberg, S. L. (2012). Fast gapped-read alignment with Bowtie 2. Nature Methods, 9(4), 357–359. 10.1038/nmeth.1923

Larsson, J., Godfrey, A. J. R., Gustafsson, P., algorithms), D. H. E. (geometric, code), E. H. (root solver, & Privé, F. (2025). eulerr: Area-Proportional Euler and Venn Diagrams with Ellipses (Version 7.0.4) [Computer software]. https://cran.r-project.org/web/packages/eulerr/index.html

Lee, S., Sorensen, J. W., Walker, R. L., Emerson, J. B., Nicol, G. W., & Hazard, C. (2022). Soil pH influences the structure of virus communities at local and global scales. Soil Biology and Biochemistry, 166, 108569. 10.1016/j.soilbio.2022.108569

Lenth, R. V., Piaskowski, J., Banfai, B., Bolker, B., Buerkner, P., Giné-Vázquez, I., Hervé, M., Jung, M., Love, J., Miguez, F., Riebl, H., & Singmann, H. (2025). emmeans: Estimated Marginal Means, aka Least-Squares Means (Version 2.0.1) [Computer software]. https://cran.r-project.org/web/packages/emmeans/index.html

Li, D., Liu, C.-M., Luo, R., Sadakane, K., & Lam, T.-W. (2015). MEGAHIT: An ultra-fast single-node solution for large and complex metagenomics assembly via succinct de Bruijn graph. *Bioinformatics (Oxford,* England), 31(10), 1674–1676. 10.1093/bioinformatics/btv033

Li, H., Handsaker, B., Wysoker, A., Fennell, T., Ruan, J., Homer, N., Marth, G., Abecasis, G., Durbin, R., & 1000 Genome Project Data Processing Subgroup. (2009). The Sequence Alignment/Map format and SAMtools. Bioinformatics (Oxford, England), 25(16), 2078–2079. 10.1093/bioinformatics/btp352

Li, Y., Adams, J., Shi, Y., Wang, H., He, J.-S., & Chu, H. (2017). Distinct Soil Microbial Communities in habitats of differing soil water balance on the Tibetan Plateau. Scientific Reports, 7, 46407. 10.1038/srep46407

Liang, J., & Karamanos, R. E. (1993). DTPA-extractable Fe, Mn, Cu and Zn. Soil Sampling and Methods of Analysis, 87–90.

Ling, N., Wang, T., & Kuzyakov, Y. (2022). Rhizosphere bacteriome structure and functions. Nature Communications, 13(1), 836. 10.1038/s41467-022-28448-9

Liu, J., Tang, Y., Bao, J., Wang, H., Peng, F., Tan, P., Chu, G., & Liu, S. (2022). A Stronger Rhizosphere Impact on the Fungal Communities Compared to the Bacterial Communities in Pecan Plantations. Frontiers in Microbiology, 13. 10.3389/fmicb.2022.899801

Love, M., Ahlmann-Eltze, C., Forbes, K., Anders, S., Huber, W., FP7, R. E., Nhgri, N., & CZI. (2023). DESeq2: Differential gene expression analysis based on the negative binomial distribution (Version 1.38.3) [Computer software]. Bioconductor version: Release (3.16). 10.18129/B9.bioc.DESeq2

Loveland, J. P., Ryan, J. N., Amy, G. L., & Harvey, R. W. (1996). The reversibility of virus attachment to mineral surfaces. *Colloids and Surfaces A: Physicochemical and Engineering Aspects*, A Collection of Papers Presented at the Symposium on Colloidal and Interfacial Phenomena in Aquatic Environments, 107, 205–221. 10.1016/0927-7757(95)03373-4

Masella, A. P., Bartram, A. K., Truszkowski, J. M., Brown, D. G., & Neufeld, J. D. (2012). PANDAseq: Paired-end assembler for illumina sequences. BMC Bioinformatics, 13(1), 31. 10.1186/1471-2105-13-31

Massicotte, P., South, A., & Hufkens, K. (2026). rnaturalearth: World Map Data from Natural Earth (Version 1.2.0) [Computer software]. https://cran.r-project.org/web/packages/rnaturalearth/index.html

Mauch-Mani, B., Baccelli, I., Luna, E., & Flors, V. (2017). Defense Priming: An Adaptive Part of Induced Resistance. Annual Review of Plant Biology, 68(1), 485–512. 10.1146/annurev-arplant-042916-041132

McLaughlin, S., Zhalnina, K., Kosina, S., Northen, T. R., & Sasse, J. (2023). The core metabolome and root exudation dynamics of three phylogenetically distinct plant species. Nature Communications, 14(1), 1649. 10.1038/s41467-023-37164-x

McMurdie, P. J., & Holmes, S. (2013). phyloseq: An R Package for Reproducible Interactive Analysis and Graphics of Microbiome Census Data. PLOS ONE, 8(4), e61217. 10.1371/journal.pone.0061217

Mehlich, A. (1984). Mehlich 3 soil test extractant: A modification of Mehlich 2 extractant. Communications in Soil Science and Plant Analysis, 15(12), 1409–1416.

Mendes, R., Garbeva, P., & Raaijmakers, J. M. (2013). The rhizosphere microbiome: Significance of plant beneficial, plant pathogenic, and human pathogenic microorganisms. FEMS Microbiology Reviews, 37(5), 634–663. 10.1111/1574-6976.12028

Mendes, R., Kruijt, M., de Bruijn, I., Dekkers, E., van der Voort, M., Schneider, J. H. M., Piceno, Y. M., DeSantis, T. Z., Andersen, G. L., Bakker, P. A. H. M., & Raaijmakers, J. M. (2011). Deciphering the rhizosphere microbiome for disease-suppressive bacteria. *Science (New York*, N.Y.), 332(6033), 1097–1100. 10.1126/science.1203980

Nicolas, A. M., Sieradzki, E. T., Pett-Ridge, J., Banfield, J. F., Taga, M. E., Firestone, M. K., & Blazewicz, S. J. (2023). A subset of viruses thrives following microbial resuscitation during rewetting of a seasonally dry California grassland soil. Nature Communications, 14(1), 5835. 10.1038/s41467-023-40835-4

Normandin, V., KotubyLJAmacher, J., & Miller, R. O. (1998). Modification of the ammonium acetate extractant for the determination of exchangeable cations in calcareous soils. Communications in Soil Science and Plant Analysis, 29(11–14), 1785–1791.

Nuzzo, A., Satpute, A., Albrecht, U., & Strauss, S. L. (2020). Impact of Soil Microbial Amendments on Tomato Rhizosphere Microbiome and Plant Growth in Field Soil. Microbial Ecology, 80(2), 398–409. 10.1007/s00248-020-01497-7

Olm, M. R., Brown, C. T., Brooks, B., & Banfield, J. F. (2017). dRep: A tool for fast and accurate genomic comparisons that enables improved genome recovery from metagenomes through de-replication. The ISME Journal, 11(12), 2864–2868. 10.1038/ismej.2017.126

Paez-Espino, D., Eloe-Fadrosh, E. A., Pavlopoulos, G. A., Thomas, A. D., Huntemann, M., Mikhailova, N., Rubin, E., Ivanova, N. N., & Kyrpides, N. C. (2016). Uncovering Earth’s virome. Nature, 536(7617), 425–430. 10.1038/nature19094

Patro, R., Duggal, G., Love, M. I., Irizarry, R. A., & Kingsford, C. (2017). Salmon provides fast and bias-aware quantification of transcript expression. Nature Methods, 14(4), 417–419. 10.1038/nmeth.4197

Pebesma, E., Bivand, R., Racine, E., Sumner, M., Cook, I., Keitt, T., Lovelace, R., Wickham, H., Ooms, J., Müller, K., Pedersen, T. L., Baston, D., & Dunnington, D. (2026). sf: Simple Features for R (Version 1.0-24) [Computer software]. https://cran.r-project.org/web/packages/sf/index.html

Pedersen, T. L. (2025). patchwork: The Composer of Plots (Version 1.3.2) [Computer software]. https://cran.r-project.org/web/packages/patchwork/index.html

Pieterse, C. M. J., Jonge, R. de, & Berendsen, R. L. (2016). The Soil-Borne Supremacy. Trends in Plant Science, 21(3), 171–173. 10.1016/j.tplants.2016.01.018

Pratama, A. A., & Elsas, J. D. van. (2018). The ‘Neglected’ Soil Virome – Potential Role and Impact. Trends in Microbiology, 26(8), 649–662. 10.1016/j.tim.2017.12.004

Quast, C., Pruesse, E., Yilmaz, P., Gerken, J., Schweer, T., Yarza, P., Peplies, J., & Glöckner, F. O. (2013). The SILVA ribosomal RNA gene database project: Improved data processing and web-based tools. Nucleic Acids Research, 41(Database issue), D590-596. 10.1093/nar/gks1219

R Core Team. (2025). R: A Language and Environment for Statistical Computing [Computer software]. R Foundation for Statistical Computing. https://www.R-project.org/

Raaijmakers, J. M., Paulitz, T. C., Steinberg, C., Alabouvette, C., & Moënne-Loccoz, Y. (2009). The rhizosphere: A playground and battlefield for soilborne pathogens and beneficial microorganisms. Plant and Soil, 321(1), 341–361. 10.1007/s11104-008-9568-6

Reinhold-Hurek, B., Bünger, W., Burbano, C. S., Sabale, M., & Hurek, T. (2015). Roots Shaping Their Microbiome: Global Hotspots for Microbial Activity. Annual Review of Phytopathology, 53(Volume 53, 2015), 403–424. 10.1146/annurev-phyto-082712-102342

Roux, S., Adriaenssens, E. M., Dutilh, B. E., Koonin, E. V., Kropinski, A. M., Krupovic, M., Kuhn, J. H., Lavigne, R., Brister, J. R., Varsani, A., Amid, C., Aziz, R. K., Bordenstein, S. R., Bork, P., Breitbart, M., Cochrane, G. R., Daly, R. A., Desnues, C., Duhaime, M. B.,…Eloe-Fadrosh, E. A. (2019). Minimum Information about an Uncultivated Virus Genome (MIUViG). Nature Biotechnology, 37(1), 29–37. 10.1038/nbt.4306

Santander, C., Aroca, R., Ruiz-Lozano, J. M., Olave, J., Cartes, P., Borie, F., & Cornejo, P. (2017). Arbuscular mycorrhiza effects on plant performance under osmotic stress. Mycorrhiza, 27(7), 639–657. 10.1007/s00572-017-0784-x

Santos-Medellín, C., Blazewicz, S. J., Pett-Ridge, J., Firestone, M. K., & Emerson, J. B. (2023). Viral but not bacterial community successional patterns reflect extreme turnover shortly after rewetting dry soils. Nature Ecology & Evolution, 7(11), 1809–1822. 10.1038/s41559-023-02207-5

Santos-Medellín, C., Estera-Molina, K., Yuan, M., Pett-Ridge, J., Firestone, M. K., & Emerson, J. B. (2022). Spatial turnover of soil viral populations and genotypes overlain by cohesive responses to moisture in grasslands. Proceedings of the National Academy of Sciences of the United States of America, 119(45), e2209132119. 10.1073/pnas.2209132119

Santos-Medellin, C., Zinke, L. A., Ter Horst, A. M., Gelardi, D. L., Parikh, S. J., & Emerson, J. B. (2021). Viromes outperform total metagenomes in revealing the spatiotemporal patterns of agricultural soil viral communities. The ISME Journal, 15(7), 1956–1970. 10.1038/s41396-021-00897-y

Scales, N. C., Huynh, K. T., Weihe, C., & Martiny, J. B. H. (2023). Desiccation induces varied responses within a soil bacterial genus. Environmental Microbiology, 25(12), 3075–3086. 10.1111/1462-2920.16494

Sims, J. T. (2000). Soil test phosphorus: Olsen P. *Methods of Phosphorus Analysis for Soils, Sediments*, Residuals, and Waters, 20.

Song, Y., Chen, D., Lu, K., Sun, Z., & Zeng, R. (2015). Enhanced tomato disease resistance primed by arbuscular mycorrhizal fungus. Frontiers in Plant Science, 6. 10.3389/fpls.2015.00786

Sorensen, J. W., Horst, A. M. ter, Zinke, L. A., & Emerson, J. B. (2023). Soil viral communities differed by management and over time in organic and conventional tomato fields (p. 2023.05.03.539301). bioRxiv. 10.1101/2023.05.03.539301

Sorensen, J. W., Zinke, L. A., Ter Horst, A. M., Santos-Medellín, C., Schroeder, A., & Emerson, J. B. (2021). DNase Treatment Improves Viral Enrichment in Agricultural Soil Viromes. mSystems, 6(5), e0061421. 10.1128/mSystems.00614-21

South, A., Michael, S., & Massicotte, P. (2024). rnaturalearthdata: World Vector Map Data from Natural Earth Used in “rnaturalearth” (Version 1.0.0) [Computer software]. https://cran.r-project.org/web/packages/rnaturalearthdata/index.html

Starr, E. P., Nuccio, E. E., Pett-Ridge, J., Banfield, J. F., & Firestone, M. K. (2019). Metatranscriptomic reconstruction reveals RNA viruses with the potential to shape carbon cycling in soil. Proceedings of the National Academy of Sciences, 116(51), 25900–25908. 10.1073/pnas.1908291116

ter Horst, A. M., Adebiyi, T. V., Hernandez, D. A., Fudyma, J. D., & Emerson, J. B. (2023). *Almond rhizosphere viral, prokaryotic, and fungal communities differed significantly among four California orchards and in comparison to bulk soil communities* (p. 2023.06.03.543555). bioRxiv. 10.1101/2023.06.03.543555

ter Horst, A. M., Fudyma, J. D., Bak, A., Hwang, M. S., Santos-Medellín, C., Stevens, K. A., Rizzo, D. M., Al Rwahnih, M., & Emerson, J. B. (2023). RNA Viral Communities Are Structured by Host Plant Phylogeny in Oak and Conifer Leaves. Phytobiomes Journal, 7(2), 288–296. 10.1094/PBIOMES-12-21-0080-R

ter Horst, A. M., Fudyma, J. D., Sones, J. L., & Emerson, J. B. (2023). Dispersal, habitat filtering, and eco-evolutionary dynamics as drivers of local and global wetland viral biogeography. The ISME Journal, 17(11), 2079–2089. 10.1038/s41396-023-01516-8

ter Horst, A. M., Santos-Medellín, C., Sorensen, J. W., Zinke, L. A., Wilson, R. M., Johnston, E. R., Trubl, G., Pett-Ridge, J., Blazewicz, S. J., Hanson, P. J., Chanton, J. P., Schadt, C. W., Kostka, J. E., & Emerson, J. B. (2021). Minnesota peat viromes reveal terrestrial and aquatic niche partitioning for local and global viral populations. Microbiome, 9(1), 233. 10.1186/s40168-021-01156-0

Trivedi, P., Leach, J. E., Tringe, S. G., Sa, T., & Singh, B. K. (2020). Plant–microbiome interactions: From community assembly to plant health. Nature Reviews Microbiology, 18(11), 607–621. 10.1038/s41579-020-0412-1

Trubl, G., Jang, H. B., Roux, S., Emerson, J. B., Solonenko, N., Vik, D. R., Solden, L., Ellenbogen, J., Runyon, A. T., Bolduc, B., Woodcroft, B. J., Saleska, S. R., Tyson, G. W., Wrighton, K. C., Sullivan, M. B., & Rich, V. I. (2018). Soil Viruses Are Underexplored Players in Ecosystem Carbon Processing. mSystems, 3(5), 10.1128/msystems.00076-18

Trubl, G., Roux, S., Solonenko, N., Li, Y.-F., Bolduc, B., Rodríguez-Ramos, J., Eloe-Fadrosh, E. A., Rich, V. I., & Sullivan, M. B. (2019). Towards optimized viral metagenomes for double-stranded and single-stranded DNA viruses from challenging soils. PeerJ, 7, e7265. 10.7717/peerj.7265

Trubl, G., Solonenko, N., Chittick, L., Solonenko, S. A., Rich, V. I., & Sullivan, M. B. (2016). Optimization of viral resuspension methods for carbon-rich soils along a permafrost thaw gradient. PeerJ, 4, e1999. 10.7717/peerj.1999

Turina, M., Ghignone, S., Astolfi, N., Silvestri, A., Bonfante, P., & Lanfranco, L. (2018). The virome of the arbuscular mycorrhizal fungus Gigaspora margarita reveals the first report of DNA fragments corresponding to replicating non-retroviral RNA viruses in fungi. Environmental Microbiology, 20(6), 2012–2025. 10.1111/1462-2920.14060

Upadhyay, S. K., Srivastava, A. K., Rajput, V. D., Chauhan, P. K., Bhojiya, A. A., Jain, D., Chaubey, G., Dwivedi, P., Sharma, B., & Minkina, T. (2022). Root Exudates: Mechanistic Insight of Plant Growth Promoting Rhizobacteria for Sustainable Crop Production. Frontiers in Microbiology, 13. 10.3389/fmicb.2022.916488

Vierheilig, H., Coughlan, A. P., Wyss, U., & Piché, Y. (1998). Ink and Vinegar, a Simple Staining Technique for Arbuscular-Mycorrhizal Fungi. Applied and Environmental Microbiology, 64(12), 5004–5007. 10.1128/AEM.64.12.5004-5007.1998

Wan, N.-F., Fu, L., Dainese, M., Kiær, L. P., Hu, Y.-Q., Xin, F., Goulson, D., Woodcock, B. A., Vanbergen, A. J., Spurgeon, D. J., Shen, S., & Scherber, C. (2025). Pesticides have negative effects on non-target organisms. Nature Communications, 16(1), 1360. 10.1038/s41467-025-56732-x

Wang, Q., Garrity, G. M., Tiedje, J. M., & Cole, J. R. (2007). Naive Bayesian classifier for rapid assignment of rRNA sequences into the new bacterial taxonomy. Applied and Environmental Microbiology, 73(16), 5261–5267. 10.1128/AEM.00062-07

White, T. J., Bruns, T., Lee, S., & Taylor, J. (1990). Amplification and direct sequencing of fungal ribosomal RNA genes for phylogenetics. PCR Protocols: A Guide to Methods and Applications, 18(1), 315–322.

Wickham, H., Chang, W., Henry, L., Pedersen, T. L., Takahashi, K., Wilke, C., Woo, K., Yutani, H., Dunnington, D., Brand, T. van den, Posit, & PBC. (2024). ggplot2: Create Elegant Data Visualisations Using the Grammar of Graphics (Version 3.5.1) [Computer software]. https://cran.r-project.org/web/packages/ggplot2/index.html

Wickham, H., Chang, W., Henry, L., Pedersen, T. L., Takahashi, K., Wilke, C., Woo, K., Yutani, H., Dunnington, D., Brand, T. van den, Posit, & PBC. (2026). ggplot2: Create Elegant Data Visualisations Using the Grammar of Graphics (Version 4.0.2) [Computer software]. https://cran.r-project.org/web/packages/ggplot2/index.html

Wickham, H., François, R., Henry, L., Müller, K., Vaughan, D., Software, P., & PBC. (2026). dplyr: A Grammar of Data Manipulation (Version 1.2.0) [Computer software]. https://cran.r-project.org/web/packages/dplyr/index.html

Wickham, H., & RStudio. (2023). tidyverse: Easily Install and Load the “Tidyverse” (Version 2.0.0) [Computer software]. https://cran.r-project.org/web/packages/tidyverse/index.html

Wickham, H., Software, P., & PBC. (2025). stringr: Simple, Consistent Wrappers for Common String Operations (Version 1.6.0) [Computer software]. https://cran.r-project.org/web/packages/stringr/index.html

Williams, A., & de Vries, F. T. (2020). Plant root exudation under drought: Implications for ecosystem functioning. New Phytologist, 225(5), 1899–1905. 10.1111/nph.16223

Williamson, K. E., Fuhrmann, J. J., Wommack, K. E., & Radosevich, M. (2017). Viruses in Soil Ecosystems: An Unknown Quantity Within an Unexplored Territory. Annual Review of Virology, 4(Volume 4, 2017), 201–219. 10.1146/annurev-virology-101416-041639

Williamson, K. E., Radosevich, M., & Wommack, K. E. (2005). Abundance and Diversity of Viruses in Six Delaware Soils. Applied and Environmental Microbiology, 71(6), 3119–3125. 10.1128/AEM.71.6.3119-3125.2005

Wu, R., Davison, M. R., Gao, Y., Nicora, C. D., Mcdermott, J. E., Burnum-Johnson, K. E., Hofmockel, K. S., & Jansson, J. K. (2021). Moisture modulates soil reservoirs of active DNA and RNA viruses. Communications Biology, 4(1), 1–11. 10.1038/s42003-021-02514-2

Wu, R., Davison, M. R., Nelson, W. C., Graham, E. B., Fansler, S. J., Farris, Y., Bell, S. L., Godinez, I., Mcdermott, J. E., Hofmockel, K. S., & Jansson, J. K. (2021). DNA Viral Diversity, Abundance, and Functional Potential Vary across Grassland Soils with a Range of Historical Moisture Regimes. mBio, 12(6), e0259521. 10.1128/mBio.02595-21

Wu, S., Shi, Z., Chen, X., Gao, J., & Wang, X. (2022). Arbuscular mycorrhizal fungi increase crop yields by improving biomass under rainfed condition: A meta-analysis. PeerJ, 10, e12861. 10.7717/peerj.12861

Xu, J., Zhang, Y., Zhang, P., Trivedi, P., Riera, N., Wang, Y., Liu, X., Fan, G., Tang, J., Coletta-Filho, H. D., Cubero, J., Deng, X., Ancona, V., Lu, Z., Zhong, B., Roper, M. C., Capote, N., Catara, V., Pietersen, G.,…Wang, N. (2018). The structure and function of the global citrus rhizosphere microbiome. Nature Communications, 9(1), 4894. 10.1038/s41467-018-07343-2

