## Supplementary figures and images for "Host community activity, but not always composition, explains viral biogeography in bulk and rhizosphere soils over a tomato growing season"

### Supplemental figure 1

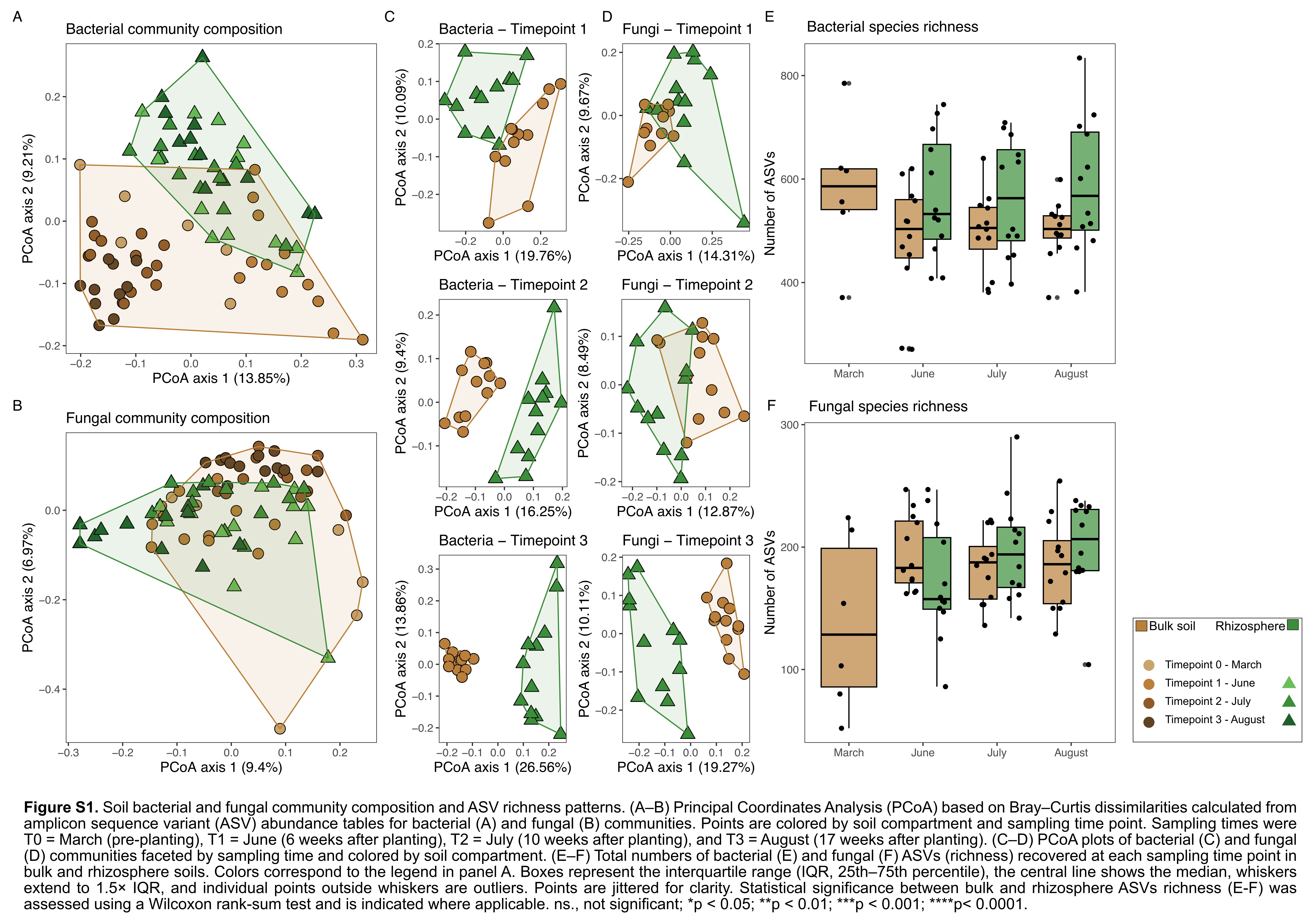

### Supplemental figure 2

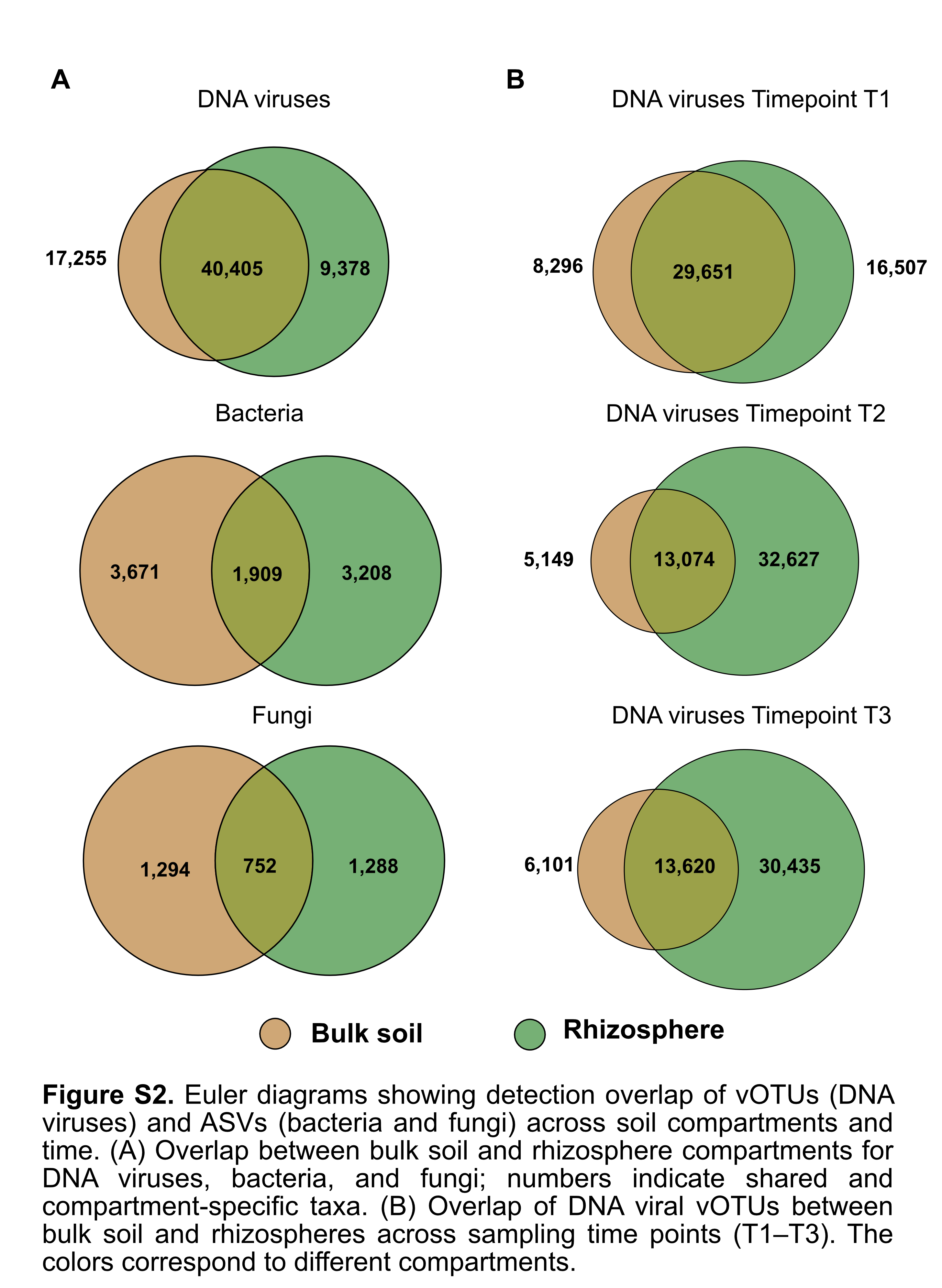

### Supplemental figure 3

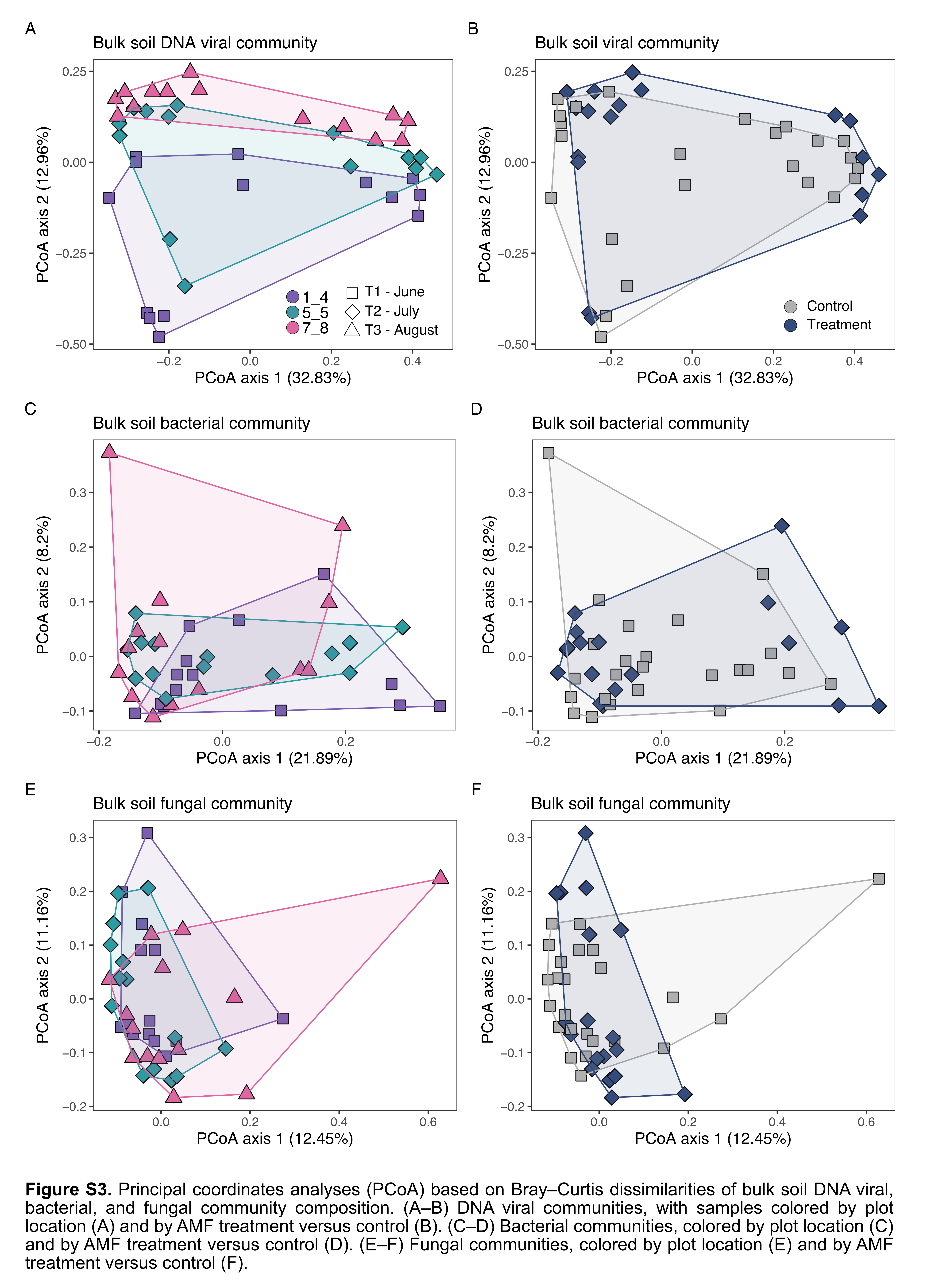

### Supplemental figure 4

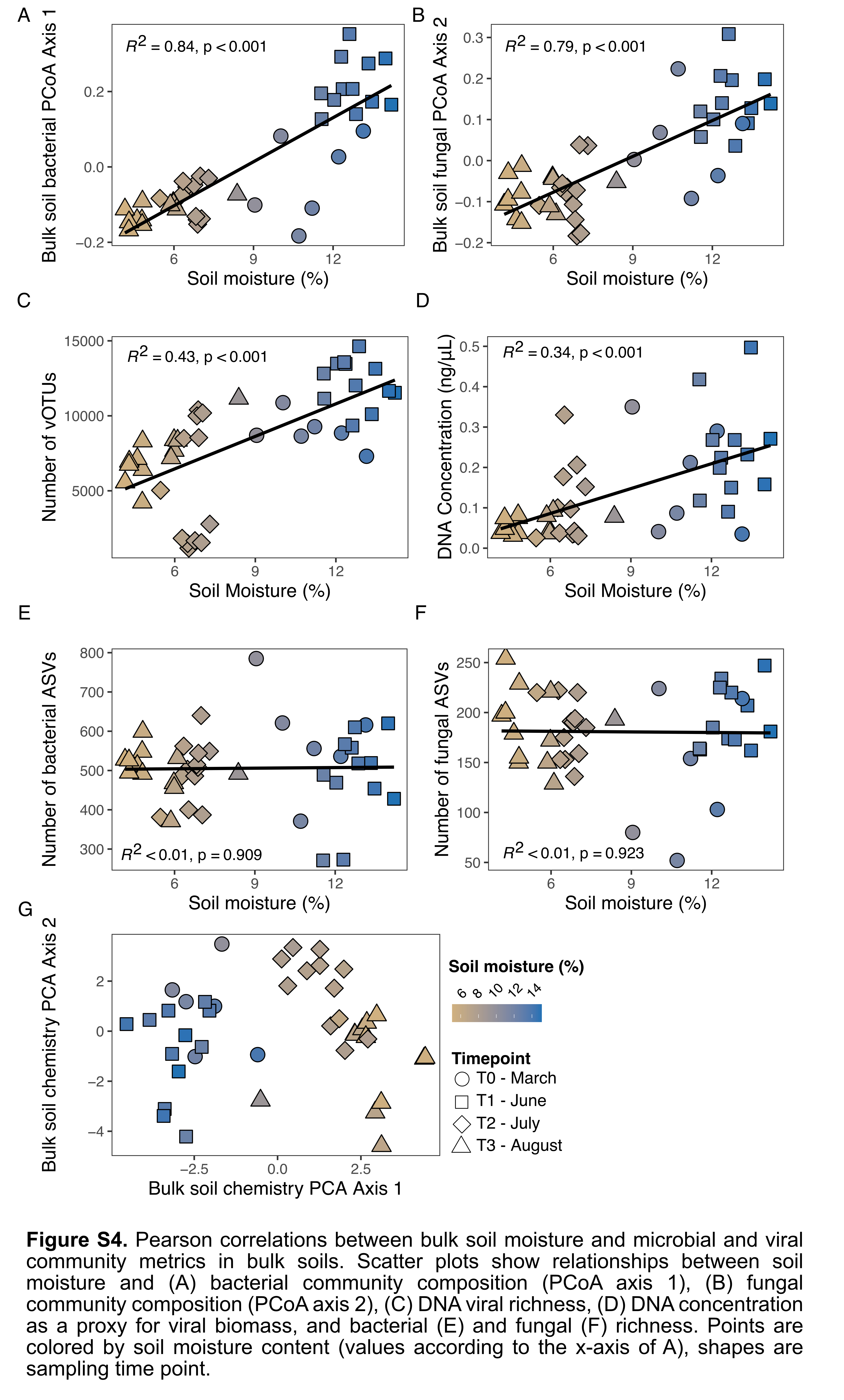

### Supplemental figure 5

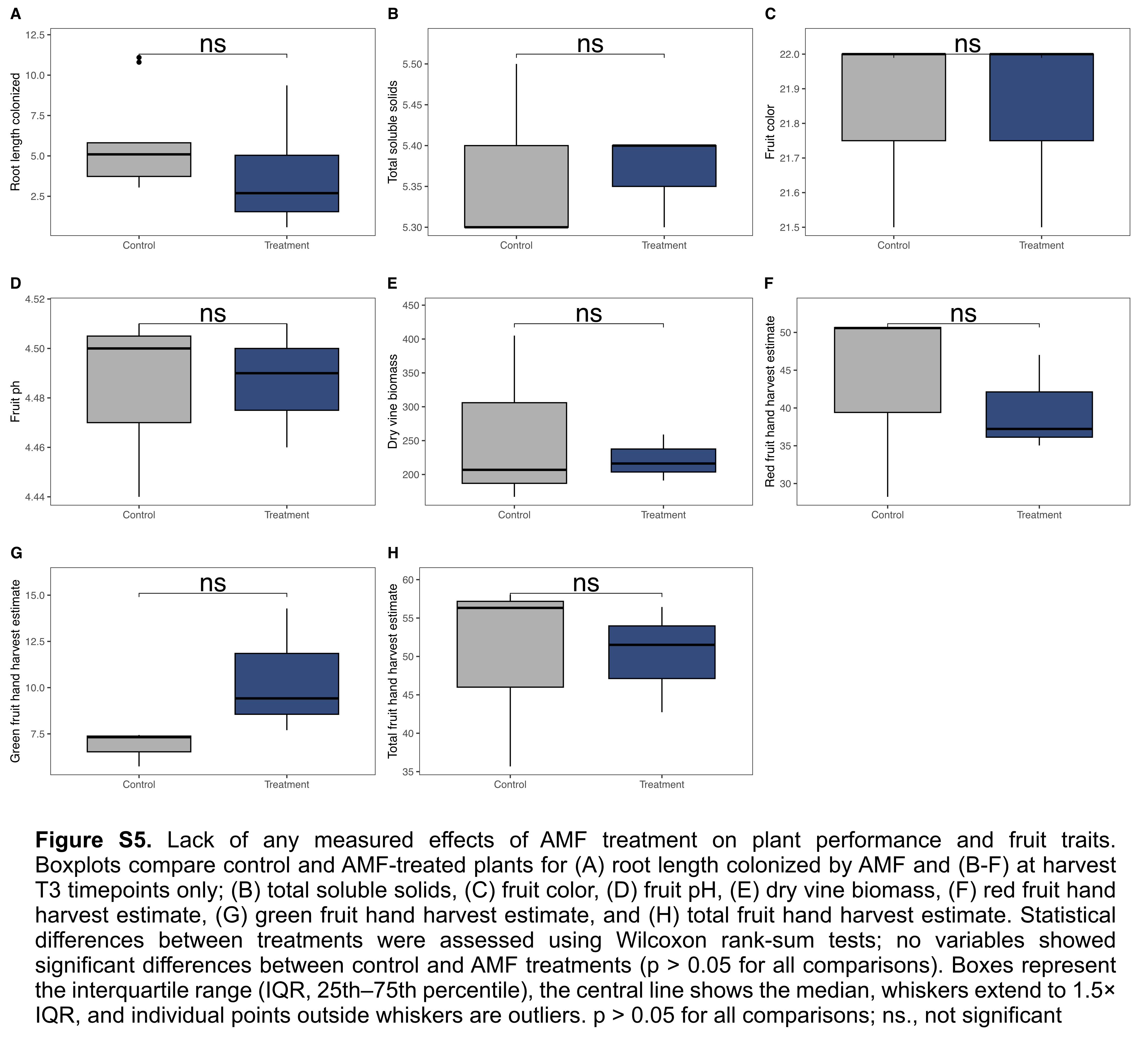
